# Production of human papillomavirus type 16 virus-like particles in Physcomitrella photobioreactors

**DOI:** 10.1101/2025.02.06.636855

**Authors:** Paul Alexander Niederau, Maria Caroline Weilguny, Sarah Chamas, Caitlin Elizabeth Turney, Juliana Parsons, Marta Rodríguez-Franco, Sebastian N.W. Hoernstein, Eva L. Decker, Henrik Toft Simonsen, Ralf Reski

## Abstract

Virus-like particles (VLPs) are self-assembling nanoparticles composed of viral structural proteins which mimic native virions but lack viral DNA and infectivity. VLPs are a resourceful class of biopharmaceuticals applied as subunit vaccines or as delivery vehicles for drugs and nucleic acids. Similar to viruses, VLPs are diverse in structure, composition, and assembly, requiring a tailored production platform aligned with the intended application. The moss plant Physcomitrella (*Physcomitrium patens*) is an emerging expression system offering humanized N-glycosylation, scalability, and adaptability to existing industry settings. Here, we used Physcomitrella to produce human papillomavirus (HPV) 16 VLPs. HPV VLPs are composed of the major structural protein L1 and are used as vaccines against HPV infections which are the main causal agent of cervical and other anogenital cancers. We characterized Physcomitrella chloroplast transit peptides, which we used for targeting of moss-produced L1 to chloroplasts, leading to higher recombinant protein yield compared to nuclear or cytoplasmic localization. We confirmed subcellular localization with confocal laser scanning microscopy and found L1 to accumulate within the chloroplast stroma. Production in 5-liter photobioreactors yielded over 0.3 mg L1 per gram fresh weight. We established a purification protocol for moss-produced L1 using a combination of ammonium sulfate precipitation and cation exchange chromatography. Purified samples were subjected to a controlled dis- and reassembly, yielding fully assembled HPV-16 L1 VLPs. This is the first report of production, purification, and assembly of VLPs in a non-vascular plant.

## Introduction

Virus-like particles (VLPs) are nanoparticles consisting of viral structural proteins (Marsian and Lomonossoff, 2016; Nooraei et al., 2021). VLPs cannot replicate and infect cells, but resemble native viruses in size, shape, and surface structure, presenting highly repetitive immunogenic epitopes. As a result, VLPs have a wide range of applications as vaccines and nanocarriers and are of high interest in biopharmaceutical research and development.

There is currently a range of VLP-based vaccines licensed and available against infectious diseases such as hepatitis B, hepatitis E, human papillomavirus (HPV), malaria, and human Norwalk virus with many more vaccine candidates in clinical trials (Nooraei et al., 2021; Gupta et al., 2023). Viral capsid proteins stimulate both the cellular and the humoral immune response, leading to the generation of cytotoxic and memory T cells as well as the production of specific antibodies *via* B cells (Ross et al., 2009; Song et al., 2011; Serradell et al., 2019; Chi et al., 2024). VLP-based vaccines offer improved safety as compared to live-attenuated or inactivated vaccines, which carry the risk of reverting to a pathogenic form (Burns et al., 2014). At the same time, VLPs don’t suffer the limitations of DNA-based vaccines, including risk of genome integration or of appropriate delivery which ultimately lowers immunogenicity (Lu et al., 2024). VLPs are not only used for their intrinsic antigenic properties but serve as nanoparticles for drug delivery by loading their inner space with cargo such as nucleic acids (Lamprecht et al., 2016; Adams et al., 2023). Besides the inner space, VLPs can also be used to display antigens on the capsid’s surface. For the treatment of pancreatic cancers, disease-unrelated VLPs were coated with respective antigens and were demonstrated to inhibit cancer growth (Cubas et al., 2011; Zhang et al., 2013). Yet, empty VLPs of Cowpea Mosaic Virus, a rod-like shaped plant virus, were shown to reduce cancer growth, despite the absence of disease-specific antigens in the vaccine (Lizotte et al., 2015). The ability of VLPs to induce strong T cell-mediated immune responses is essential in VLP-based cancer vaccines (Nooraei et al., 2021). Overall, VLPs possess characteristics that make them highly versatile for applications in vaccine development, targeted drug delivery, gene therapy, and immunotherapy, offering significant potential in modern biomedicine.

VLPs are classified based on various structural characteristics, one of which is the presence or absence of a lipid envelope (Nooraei et al., 2021). In enveloped VLPs (eVLPs) the viral capsid proteins are imbedded within a lipid membrane acquired during particle assembly, typically from intracellular membranes in the secretory pathway or the outer plasma membrane. Additional structural characteristics include the number and types of capsid proteins, as well as their organization into single, double, or triple-layered architectures. In some instances, capsid proteins require proteolytic processing or N-glycosylation to enable budding of assembled particles from the host cell’s membrane and acquisition of the lipid envelope (Welsch et al., 2007; Zlotnick and Mukhopadhyay, 2011; Kim et al., 2014). Furthermore, viral antigens displayed on the surface of the lipid membrane are often glycosylated, which significantly contributes to the immunogenicity of eVLP-based vaccines (Mortola and Roy, 2004). One example is the major glycoprotein of Ebola virus for which the distinct glycosylation pattern is crucial for the formation of neutralizing antibodies (Peng et al., 2022). Other notable cases include hepatitis C virus and Zika virus – both of which currently lack a licensed vaccine – where the induction of neutralizing antibodies depends on viral glycoproteins (Torresi, 2017; Hasan et al., 2018). Consequently, the choice of VLP, its structural complexity, and intended application play a crucial role in determining the appropriate expression system and guiding cell line development.

The moss Physcomitrella has emerged as an alternative for biopharmaceutical production, offering several advantages over traditional systems (Reski et al., 2015; Reski, 2018; Decker and Reski, 2020). Its haploid dominant phase and high rate of homologous recombination enable precise gene targeting (Strepp et al., 1998; Wiedemann et al., 2018; Rempfer et al., 2022). This feature facilitated the modification of the N-glycosylation pathway, eliminating plant-specific modifications, such as β1,2- xylosylation, α1,3-fucosylation, and Lewis A epitopes (Koprivova et al., 2004; Parsons et al., 2012; Decker et al., 2014). Moreover, enzymes necessary for sialic acid synthesis, activation, and linkage to protein N-glycans were introduced (Bohlender et al., 2020; Bohlender et al., 2022). These modifications have resulted in more homogeneous and human-like N-glycan structures on recombinant proteins. The potential of Physcomitrella as a biopharmaceutical production platform is demonstrated by several products in development. For example, moss-produced α-galactosidase A has successfully completed clinical phase Ib (Shen et al., 2016; Hennermann et al., 2019). Other moss-produced proteins include human factor H (FH) and synthetic FH-related multitarget regulators MFHR1 and MFHR13, which are potential treatments for complement-related disorders (Büttner-Mainik et al., 2011; Michelfelder et al., 2017, 2018; Top et al., 2019; Ruiz-Molina et al., 2022a) such as age-dependent macular degeneration (Hector et al., 2025). Factor H produced in other systems faced difficulties in its application, such as *Pichia pastoris*-produced factor H exhibiting non-optimal glycosylation and reduced half-life (Schmidt et al., 2011; Kerr et al., 2021). Importantly, Physcomitrella cultivation in photobioreactors is tightly controlled (Hohe et al., 2002), and thus compliant with Good Manufacturing Practice (GMP) standards, ensuring high-quality production of recombinant proteins (Reski et al., 2018; Decker and Reski, 2020). As opposed to transiently transfected *Nicotiana benthamiana* plants cultivated in green houses, cultivation of stable transgenic Physcomitrella cell lines in bioreactors is more in line with current production processes applied in industry, thereby facilitating adaption to industry settings (Benvenuto et al., 2023). In addition, cultivation of plant material in bioreactors allows greater scalability, which is a relevant aspect in transitioning to industrial scale production. The company Medicago, focusing on VLP production in transiently transfected *N. benthamiana*, announced challenges in transitioning with their platform to industrial scale production as a reason for seizing further operations in 2023 (Mitsubishi Chemical Group Corporation, 2023). In contrast, the company Eleva upscaled protein production in Physcomitrella from 200 L single-use bioreactors (Niederkrüger et al., 2019) to 1000 L photobioreactors (Eleva, 2025). In summary, Physcomitrella is an ideal production host for complex biopharmaceuticals, including viral antigens or glycosylated proteins, with great prospects regarding scalability and transition to industrial production (Rosales-Mendoza et al., 2014). The first moss-produced vaccine candidate, an ENV-derived multi-epitope HIV chimeric protein, was immunogenic in mice (Orellana-Escobedo et al., 2015). However, until now, there was no record of VLP-production in Physcomitrella, or any other non-vascular plant.

VLPs of the human papillomavirus (HPV) are of great medicinal relevance due to their use as vaccines to prevent HPV infection. HPV infections are a major source of cancer, causing approximately 690,000 cases annually (zur Hausen et al., 1981; zur Hausen, 2002; de Martel et al., 2020). The most prevalent type is cervical cancer, accounting for about 80% of HPV-attributable cancers. Other HPV-related cancers include anogenital and oropharyngeal cancers (Wu et al., 2024). HPV belongs to the *Papillomaviridae*, comprising nearly 200 different types, which are commonly transmitted sexually (Ljubojevic and Skerlev, 2014). The majority of HPV-attributable cancers are linked to two types in particular, namely HPV-16 and HPV-18 (de Martel et al., 2020; Wu et al., 2024). HPV is a non-enveloped double stranded DNA virus with a genome size of 8000 bp, encoding six early regulatory proteins (E1, E2, E4 – E7) and two late structural proteins (L1, L2) (Roden and Stern, 2018). L1 proteins multimerize into pentamers, the capsomers, and L1 capsomers and L2 monomers are imported into the nucleus *via* nuclear localization signals (NLS), where HPV virions assemble (Cerqueira and Schiller, 2017). The final HPV-16 virion is composed of 360 L1 monomers, organized in 72 pentamers, and 72 L2 monomers and is reinforced by inter-pentameric disulfide bonds (Baker et al., 1991; Buck et al., 2005). While the role of L2 in virion assembly differs from type to type, L1 can self-assemble into VLPs in the absence of L2 (Kirnbauer et al., 1992). HPV L1 VLPs are mono-layered and do not contain a lipid envelope. Because the viral life cycle depends on differentiated epithelial cells, it is difficult to obtain large quantities of virions. Therefore, current HPV vaccines are based on L1 VLPs. Although VLP-based HPV vaccines are currently produced in *Escherichia coli*, *Saccharomyces cerevisiae*, and baculovirus-infected insect cells (World Health Organization, 2024), plant-based expression systems are gaining attention in this respect (Scotti and Rybicki, 2013; Rybicki, 2020).

While first attempts to produce L1-based VLPs in plants had low yields (Biemelt et al., 2003; Varsani et al., 2003), considerable improvements have been achieved, e.g. by codon optimization and subcellular targeting strategies in transiently transformed tobacco (Maclean et al., 2007). A human codon-optimized version (62% GC content) resulted in highest yields as compared to *N. benthamiana* optimized (49% GC) and the native sequence (37% GC), presumably not due to the GC content but due to inhibitory 5’ sequences, which act on transcription and mRNA processing (Maclean et al., 2007; Hitzeroth et al., 2018). Further, these authors found chloroplast-targeted L1 yield highest at 0.5 mg / g fresh weight (FW) followed by cytoplasmic L1 (without the C-terminal 22 aa NLS), whereas L1 targeted to the ER was not detectable. Similarly, Zahin et al. (2016) reported 0.25 mg L1 / g FW *via* chloroplast targeting while leaving the NLS intact. Although *N. benthamiana* remains a prominent option, other plants are increasingly explored for the efficient production of biopharmaceuticals.

Here, we analyzed the expression of HPV-16 L1 in Physcomitrella. We characterized five Physcomitrella chloroplast transit peptides and investigated the effect of subcellular L1 localization on recombinant protein yields. We further established L1 production in photobioreactors and subsequent purification of the product. Finally, the assembly of moss-produced HPV-16 L1 to VLPs was demonstrated with transmission electron microscopy. Our goal was to obtain proof-of-concept to produce a simple but medicinally relevant VLP before progressing towards more complex targets such as eVLPs consisting of multiple proteins and layers.

## Materials and methods

### Plant material

Physcomitrella (new species name: *Physcomitrium patens* (Hedw.) Mitt.; IMSC accession number 41269), was cultivated according to Reski and Abel (1985) and Decker et al. (2015).

### Optimization of HPV-L16 L1 coding sequence

The coding sequence (CDS) of h(human codon-optimized)L1 (GenBank acc. no. DQ067889, Maclean et al., 2007) was used for optimization using physCO (https://www.plant-biotech.uni-freiburg.de/tools/physco/; Top et al., 2021). Putative alternative splicing motifs AGGT were identified *via* search function. MicroRNA-binding sites were predicted *via* psRNATarget with the library of 280 published Physcomitrella microRNAs and the default settings *Schema V2* (2017 release) (http://plantgrn.noble.org/psRNATarget/; Dai and Zhao, 2011; Dai et al., 2018). Alternative splice sites and microRNA-binding sites were removed and substituted by synonymous codons based on http://www.kazusa.or.jp/codon/. Sequence editing was done in Benchling (https://www.benchling.com/).

### Expression vectors

Chloroplast transit peptides (CTPs) were selected from a list of proteins with experimentally confirmed cleavage of a predicted CTP (Hoernstein et al., 2024). CTP-eGFP vectors were constructed based on a vector containing an eGFP CDS with an N-terminal linker (sequence: GGGGGA) under control of the PpActin5 promoter (Weise et al., 2006) and nos terminator (Hoernstein et al., 2023). CTP candidates were amplified from Physcomitrella cDNA and inserted at the N-terminus of the linker using Gibson Assembly (Gibson et al., 2009). The Citrine-L1 fusion constructs are based on a vector containing the PpActin5 promoter, nos terminator and citrine CDS (Wiedemann et al., 2018). The CDS of pL1, pL1Δ22 and CTP-pL1 were amplified using Phusion^TM^ polymerase (Thermo Fisher Scientific, Waltham, USA), cloned into the pJet1.2 backbone (Thermo Fisher Scientific) and ligated into the expression vector using T4 Ligase (Thermo Fisher Scientific), *via Xho*I and *BamH*I sites. The pL1 expression vectors were constructed based on a vector containing the PpActin5 promoter, nos terminator and hpt cassette (Ruiz-Molina et al., 2022a). The pL1 sequence was synthesized by Genewiz (Leipzig, Germany). For constructs pL1 and pL1Δ22, the pL1 CDS was amplified using Phusion^TM^ polymerase (Thermo Fisher Scientific) and cloned as described above. For construct CTP-pL1, the pL1 and CTPc5 CDS were amplified and inserted into the backbone using Gibson Assembly. Vectors were sequenced by Eurofins Genomics (Ebersberg, Germany). Vector maps are compiled in **Supplementary Figure S1**. Primers were synthesized by Eurofins Genomics and are compiled in **Supplementary Table S1**.

### Protoplast preparation and transformation

Physcomitrella protoplasts were prepared and transformed based on existing protocols (Hohe and Reski, 2002; Schween et al., 2003; Hohe et al., 2004; Decker et al., 2015). Transformed protoplasts were incubated at 22 °C for 24 h in the dark and subsequently for 72 h at a 16/8-h light/dark photoperiod and light intensity of 50–70 μmol m^-2^ s^-1^ before plating on KnopME + 25 mg/L hygromycin agar plates covered with cellophane for two weeks. Surviving colonies were transferred to non-selective conditions for two weeks before a second selection on KnopME + 50 mg/L hygromycin.

### Fluorescence microscopy

Protoplasts were analyzed 2-3 days after transfection with an AxioCam MRc5 camera on an Axioplan2 stereo microscope (Zeiss, Oberkochen, Germany). Pictures were taken with a 40x objective or a 63x objective with immersion oil. A FITC filter was used for eGFP, a rhodamine filter for chlorophyll autofluorescence, and a FITC_LP filter for the combination of the two. Images were edited with GIMP 2.10.22 and ImageJ. For confocal microscopy, an AXIO Observer.Z1 Inverted Fluorescence Motorized Phase Contrast Microscope Pred 7 with the objective Plan-Apochromat 63x/1.4 Oil DIC M27 (Zeiss) was used. EGFP signals were detected at 488 / 509 nm, and chlorophyll autofluorescence at 587 / 610 nm. Images were edited in ZEN 3.5 (Blue edition, ZEN Lite), ImageJ, and GIMP 2.10.22.

For analysis of subcellular targeting, an inverted confocal laser scanning microscope LSM 880 with the objective LD LCI Plan-Apochromat 40x/1.2 H_2_O autocorr and FastAiryScan mode (Zeiss) was used. Citrine signals were detected at 488 / 516 nm, and chlorophyll autofluorescence at 561 / 595 nm. Images were edited in ZEN 3.9 (Blue edition) and processed *via* deconvolution using default setting and orthogonal projection using maximum intensity projection.

### Biomass accumulation

Flasks containing 30 mL KnopME medium were inoculated at a density of 60 mg dry weight (DW)/L and cultured for 4 weeks at the conditions described above. Biomass DW was measured by filtering 3x 10 mL of culture through miracloth and subsequent drying at 105 °C for 2 hours. Biomass accumulation was analyzed by linear regression analysis. Slopes were compared using ANCOVA and post hoc multiple comparisons test (TukeyHSD). The analysis was conducted in RStudio (version 4.2.2) and GraphPad Prism (version 8.1).

### Bioreactor operation

Stirred tank bioreactors (Getinge, Sweden) were inoculated at 60 mg DW/L moss density in 5 L KnopME. The light intensity was set to 160 µmol m^-2^ s^-1^ and increased to 350 μmol m^-2^ s^-1^ after 4 days. At day 4, auxin 1-Naphtalene acetic acid (NAA, Sigma, St. Louis, USA) dissolved in 0.5 M KOH was added to a final concentration of 5 µM. Throughout the bioreactor run, the pH was kept at pH 5.8, temperature at 22 °C, and aeration at 0.3 vvm (volume air per volume media per minute) with 2% CO_2_. The culture was continuously agitated with a pitched 3 blade impeller at 500 rpm.

### Protein extraction, purification and VLP assembly

For small scale extraction, 30 – 60 mg fresh weight (FW) of vacuum-filtered moss material was combined with a glass and a steel bead (each 3 mm in diameter) in a 2 mL reaction tube, frozen in liquid nitrogen and the tissue lysed using a tissue lyser for 2 min at 28 Hz. Homogenized material was resuspended in PBS (2.7 mM KCl, 10 mM Na_2_HPO_4_, 1.8 mM KH_2_PO_4_, 0.137 M NaCl, pH 7.2), 0.01% (v/v) Tween20, and protease inhibitor (P9599, Sigma) in a ratio 3 mL/g. Samples were placed in an ice-cold sonication bath for 15 min, centrifuged for 30 min at 14,000 x g and 4 °C, and the supernatant taken off for further analysis.

For large scale protein extraction, 5 g FW of vacuum-filtered moss material were frozen in liquid nitrogen and ground to a fine powder. The material was combined with 22 mL PBS and 0.01% (v/v) Tween20, 50 mM DTT and 100 µL protease inhibitor. The mixture was homogenized using an ULTRA-TURRAX (IKA, Staufen, Germany) at 10,000 rpm for 10 min on ice. Next, the extract was treated with a Q500 sonicator using a CL-334 converter (QSonica, Newtown, USA) at amplitude 55%, 10s on & 40s off for 20 min at 8 °C. Debris was removed by two centrifugations at 4500 x g for 10 min and 20,000 x g for 20 min, both at 4 °C. The sample was combined with solid (NH_4_)_2_SO_4_ to a final concentration of 45% (w/v) and proteins precipitated under stirring for 1 h at 8 °C. Amounts of (NH_4_)_2_SO_4_ were calculated using an online tool (https://files.encorbio.com/protocols/AM-SO4.htm) (EnCor Biotechnology Inc., Gainesville, USA). Precipitated proteins were pelleted by centrifugation at 12,000 x g for 10 min at 8 °C. Pellets were dissolved in 4 mL cation exchange (CaEx) chromatography binding buffer (2.7 mM KCl, 10 mM Na_2_HPO_4_, 1.8 mM KH_2_PO_4_, 0.068 M NaCl, 0.01% (v/v) Tween 20, 5% (v/v) Glycerol, 5 mM DTT, pH 6.8) and dialyzed (20k MW cut off, 88405, Thermo Fisher Scientific) overnight against the same buffer at 8 °C. The sample was diluted to a final volume of 10 mL using binding buffer, and passed through a 0.22 µm filter (Roth, Karlsruhe, Germany) before loading onto a 1 mL HiTrap SP HP cation exchange column (Cytiva, Marlborough, USA), connected to an ÄKTA^TM^ start (Cytiva). The column was washed with 5 column volumes (CV) of 0.3 M NaCl (based on binding and elution buffer) and L1 eluted *via* 4 CV of elution buffer (2.7 mM KCl, 10 mM Na_2_HPO_4_, 1.8 mM KH_2_PO_4_, 1 M NaCl, 0.01% (v/v) Tween 20, 5% (v/v) Glycerol, 5 mM DTT, pH 6.8)

Eluted fractions containing L1 were pooled and dialyzed against disassembly buffer (22.7 mM KCl, 10 mM Na_2_HPO_4_, 1.8 mM KH_2_PO_4_, 0.068 M NaCl, 0.05% (v/v) Tween20, 5 mM DTT, pH 8.5) overnight at 8°C with 3x buffer exchange. Samples were concentrated 5x using a vacuum concentrator (Concentrator plus, Eppendorf, Hamburg, Germany). For reassembly, samples were dialyzed against reassembly buffer (22.7 mM KCl, 10 mM Na_2_HPO_4_, 1.8 mM KH_2_PO_4_, 1 M NaCl, 0.05% (v/v) Tween20, pH 6) overnight at 8 °C with 3x buffer exchange.

### SDS-PAGE and western blot

SDS-PAGE and western blots were performed based on Bohlender et al. (2022). Two systems were used for protein separation, either 7.5% polyacrylamide gels (Mini-PROTEAN® TGX^TM^ Precast Gels, Rio- Rad, Munich, Germany) in TGS buffer at 120 V, or 4 – 12% Bis-Tris gels (mPAGE^TM^ Precast Gels, Merck, Darmstadt, Germany) in MOPS buffer at 150 V. As first antibody served 1:5000 diluted CamVir1 (Thermo Fisher Scientific), as second antibody peroxidase-linked rabbit anti-mouse secondary antibody (Cytiva) diluted 1:12,500. For dot blots, protein samples were blotted on a PVDF membrane (Cytiva) using a filtration manifold kit (SRC-96/1, Schleicher&Schuell, Dassel, Germany). Antibody H16.V5 (AntibodySystems, Schiltigheim, France) was diluted 1:5000. Dot blots were developed as described for western blots. Yeast-produced HPV-16 L1 (Abcam, Cambridge, UK) was used as a positive control.

### L1 quantification

The ELISA protocol was based on Studentsov et al. (2002). In short, 96-well microtiter plates (Greiner, Kremsmünster, Austria) were coated with 1:500 CamVir1 antibody (Thermo Fisher Scientific) in PBS overnight at 8 °C. Wells were washed twice with washing buffer (1x PBS, 0.05% (v/v) Tween20) and blocked with blocking buffer (1x PBS, 1% (w/v) polyvinyl alcohol MW 47,000, 0.05% (v/v) Tween20) for 3 h at 37 °C. After 3x washing, samples were diluted in blocking buffer and incubated in the wells for 1.5 h at 37 °C. Wells were washed 3x before adding 1:1000 diluted polyclonal anti-HPV-16 L1 antibody from rabbit (Cusabio, Houston, USA) for 1 h at 37 °C. After 5x washing, wells were incubated with 1:1000 diluted anti-rabbit HRP-conjugated antibody NA934 (Cytiva) for 1 h at 37°C. Wells were washed 3x before adding 100 µL TMB substrate (Thermo Fisher Scientific). The reaction was stopped with 50 µL of 2 M H_2_SO_4_. Absorbance was measured at 450 / 595 nm for reference. The signal of a wildtype sample was subtracted to account for background signal.

### Transmission electron microscopy

The TEM protocol was based on Abel et al. (1989). Negative staining of purified VLPs and moss extracts was performed using 300 Mesh glow discharged formvar/carbon-coated copper grids (Electron Microscopy Sciences, Hatfield, USA). Five µL of sample were applied on a grid and incubated for 5 min at RT. Grids were washed 3x with water and stained with 2% (w/v) uranyl acetate. After 30 sec, uranyl acetate was removed with a filter paper, grids were air dried, and inspected with a Hitachi HT7800 TEM (Tokyo, Japan) coupled to a Xarosa CMOS camera (Emsis, Münster, Germany).

## Results

### Adaptation of L1 coding sequence for Physcomitrella-optimized expression

The coding sequence (CDS) of HPV-16 L1 was optimized for expression in Physcomitrella. The goal was to adapt the CDS regarding codon usage and preventing potential degradation through microRNAs and splicing. A human codon-optimized version (hL1, **Supplementary Figure S2**) was used as a basis for optimization since it had led to higher yields than plant-optimized versions or the native viral sequence in *Nicotiana benthamiana* (Maclean et al., 2007).

For codon optimization, physCO (Top et al., 2021) was used, which changed nine out of 505 codons and slightly increased the GC content from 62.25% to 62.85% (**Supplementary Figure S2**). Next, the CDS was scanned for the motif AGGT, which is the most common motif at exon-intron borders for alternative splicing in Physcomitrella (Top et al., 2021). Transformation of a heterologous cDNA containing the AGGT motif led to alternative splicing in Physcomitrella, a process referred to as heterosplicing (Top et al., 2021). Scanning the physCO-optimized hL1 sequence yielded 5 AGGT-motifs which were replaced by synonymous codons (**Supplementary Figure S2**). MicroRNAs bind complementary sequences in the mRNA and thereby cause inhibition of translation or degradation of mRNA (Jones-Rhoades et al., 2006; Khraiwesh et al., 2010). One microRNA binding site was predicted within the CDS using psRNATarget (Dai and Zhao, 2011; Dai et al., 2018). Therefore, we changed three bases within the predicted binding region using synonymous codon substitutions. The final Physcomitrella-optimized CDS was named pL1 (**Supplementary Figure S2**) and differs from hL1 by 17 out of 1515 nucleotides without changing the amino acid (aa) sequence.

### Characterization of chloroplast transit peptides

Different constructs were designed to target L1 to specific cellular compartments, including chloroplasts. Fusing chloroplast transit peptides (CTPs) to the L1 CDS led to increased protein yields in tobacco as compared to constructs lacking CTPs (Maclean et al., 2007; Zahin et al., 2016). To assess whether this is also true for L1 production in Physcomitrella, a selection of Physcomitrella-derived CTPs was characterized.

Initially, the CTP of FtsZ1-1 (Gremillon et al., 2007) was considered. However, cleavage site analysis *via* TargetP-2.0 (Armenteros et al., 2019) predicted 37 aa of this CTP to remain at the N-terminus of L1 after chloroplast import. This CTP was rejected because the additional aa could change L1 properties and hamper VLP assembly. A list of alternative CTPs was derived from the Physcomitrella N-terminome (Hoernstein et al., 2024) (**Supplementary Table S2**). Here, experimentally detected protein N-termini were compared with TargetP-2.0 predictions for cleavable N-terminal targeting sequences. Five CTPs were selected where an exact match between prediction and experimentally validated N-terminus was observed across different experiments (Hoernstein al., 2024), namely CTPc5, CTPc10.1, CTPc10.2, CTPc21, CTPc22 (**Table 1**). Their sequences were fused *in silico* to the N-terminus of L1 and analyzed with TargetP-2.0 to validate the correct cleavage of the fusion.

**Table 1.** ***In-silico* analysis of Physcomitrella-derived chloroplast transit peptides.** CTPs were selected based on mass spectrometry datasets (Hoernstein et al., 2024). *In-silico* predictions of likelihood of chloroplast localization target sequence cleavage were performed with TargetP-2.0.

| Name of CTP | Originating from protein model | Predicted cleavage of CTP-L1 | Length (aa) | Likelihood of GFP chloroplast localization | Likelihood of L1 chloroplast localization |
| --- | --- | --- | --- | --- | --- |
| FtsZ1-1 | Pp3c22_4940V3.1 | 37 aa remain on N-terminus | 84 | 0.95 | 0.92 |
| CTPc5 | Pp3c5_1370V3.1 | - | 74 | 0.987 | 0.943 |
| CTPc10.1 | Pp3c10_10490V3.1 | - | 63 | 0.456 | 0.229 |
| CTPc10.2 | Pp3c10_2940V3.1 | - | 70 | 0.908 | 0.845 |
| CTPc21 | Pp3c21_9980V3.1 | - | 43 | 0.956 | 0.891 |
| CTPc22 | Pp3c22_5470V3.1 | - | 62 | 0.939 | 0.739 |
| CTPc22noMet | Pp3c22_5470V3.1 | - | 61 | 0.913 | 0.727 |

Our *in-silico* analysis indicated CTPc22 to end with a methionine, which could introduce an alternative translation initiation site, resulting in translation of L1 without the CTP (**Table 1**). Therefore, an alternative version lacking the C-terminal methionine (CTPc22noMet) was included. The likelihoods for chloroplast localization were above 70% for all candidates except CTPc10.1 (**Table 1**), which was rejected.

The remaining five CTPs (CTPc5, CTPc10.2, CTPc21, CTPc22, CTPc22noMet) were fused to the CDS of eGFP (**Supplementary Figure S1**) and expressed in Physcomitrella protoplasts. Analysis *via* fluorescence microscopy confirmed localization of eGFP signals in chloroplasts for CTPc5, CTPc10.2, CTPc22, and CTPc22noMet, but not for CTPc21 (**Supplementary Figure S3**). Of the four remaining CTPs, CTPc5 had the highest likelihood for plastid localization (**Table 1**) and were examined *via* confocal microscopy (**Supplementary Figure S3**). Finally, we selected CTPc5 for the chloroplast-targeting of L1.

### Targeting of L1 to nuclei and chloroplasts

To evaluate the impact of L1 localization on product yield, we generated three variants (pL1, pL1Δ22, CTP-pL1). pL1 contains the full-length pL1 CDS, which includes the native HPV-16 NLS (Cerqueira and Schiller, 2017). For pL1Δ22 the NLS was removed. In CTP-pL1 the Physcomitrella CTPc5 was fused to the 5’ of the pL1 CDS, including the NLS.

After generation of these CDS variants, we aimed to evaluate L1 targeting to subcellular compartments. The CDS of pL1, pL1Δ22 and CTP-pL1 were fused to the 5’ end of Citrine CDS (**Supplementary Figure S1**). Physcomitrella protoplasts were transfected with the plasmids and Citrine localization was analyzed using fluorescence microscopy. A vector encoding Citrine without any L1 variant fusion served as a control.

Cells transformed with the control vector exhibited predominantly cytoplasmic Citrine signals, whereas diffuse signals also occurred in nuclei (**Figure 1**). For pL1-Citrine, the signal localized to nuclei, demonstrating that the HPV-16 NLS is functional in Physcomitrella. This is further supported by the fact that removal of the NLS in pL1Δ22-Citrine results in cytoplasmic localization of the signal. For CTP-pL1-Citrine, the signal is predominantly in chloroplasts, but is not evenly distributed, as seen for CTP-eGFP fusions lacking L1 (**Supplementary Figure S3**). Instead, the pL1-Citrine fusion concentrates in several spots of high fluorescence intensity and less than 1 µm in diameter. It is unclear whether this difference is due to distinct features of GFP and Citrine or due to the presence of L1 and potential formation of L1 multimeric structures. Overall, we confirmed successful L1 targeting to the intended compartments.

**Figure 1.**
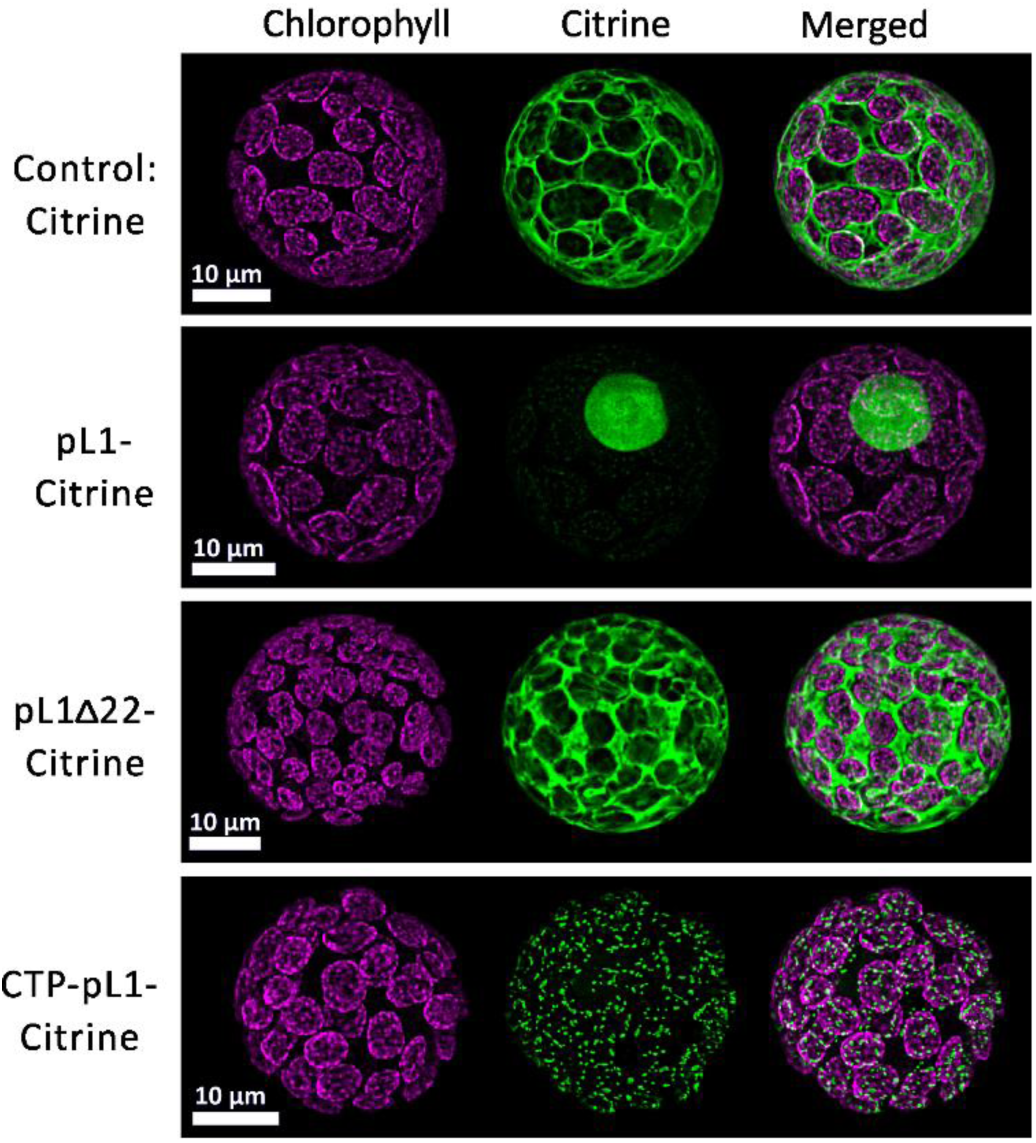
Subcellular localization of L1 variants. Confocal microscopy images of Physcomitrella protoplasts transformed with constructs encoding (CTP-)pL1(Δ22)-Citrine fusion proteins or Citrine alone. For each sample, chlorophyll autofluorescence in magenta is seen on the left, Citrine signals in the middle and the merged signals on the right. In the control and pL1Δ22-Citrine, the Citrine signals do not enter the chloroplasts and are mainly in the cytoplasm. For pL1-Citrine, the Citrine signals are in the nucleus. In the CTP-pL1-Citrine sample, the Citrine signals form small spots of high intensity in the chloroplasts.

### Characterization of L1 production

After validating the subcellular localization, pL1, pL1Δ22 and CTP-pL1 expression constructs were cloned (**Supplementary Figure S1**). The linearized constructs were transferred into Physcomitrella protoplasts and obtained plants were screened for productivity by western blot and ELISA.

Western blot analysis revealed the L1 monomer of 56 kDa (pL1, CTP-pL1) and 54 kDa (pL1Δ22) (**Figure 2**). In addition, pL1 and CTP-pL1 lines exhibited a band at approximately 54 kDa. All lines showed another band at around 45 kDa, even though the signal was relatively weak in CTP lines and more prominent in pL1Δ22 lines. Smaller protein isoforms can originate from degradation or DNA or RNA, resulting in truncated transcripts and, subsequently, truncated protein isoforms (Murén et al., 2009; Top et al., 2021). The latter possibilities were ruled out by amplifying the L1 CDS *via* PCR from cDNA, which yielded amplicons of the expected sizes, corresponding to the western blot bands of 56 kDa (pL1, CTP- pL1) and 54 kDa (pL1Δ22), but not additional amplicons (**Supplementary Figure S4**). Also, two bands are visible at 130 – 180 kDa, of which the highest band at 180 kDa is also present in the yeast-produced L1 control. This pattern of two close-by bands repeats another two times at higher molecular weight outside of the marker range (**Figure 2**). Addition of reducing agents DTT or ascorbic acid to the extraction buffer markedly reduces the western blot signal intensity above 180 kDa (**Supplementary Figure S5**). We hypothesize that in the absence of reducing agents, reactive oxygen species (ROS) in the plant extract cross-link L1 with itself and/or host proteins. Irrespective of the presence or absence of reducing agents during extraction and purification, two prominent bands in the range of 130 and 180 kDa are present in all western blots (**Figures 2 and 4, Supplementary Figure S7**). Western blot and ELISA analysis indicate higher yields in CTP-pL1 lines as compared to pL1 or pL1Δ22 lines. In shaking flasks, the best producer is CTP-pL1 30 with 142 µg L1/g FW (**Figure 2**).

**Figure 2:**
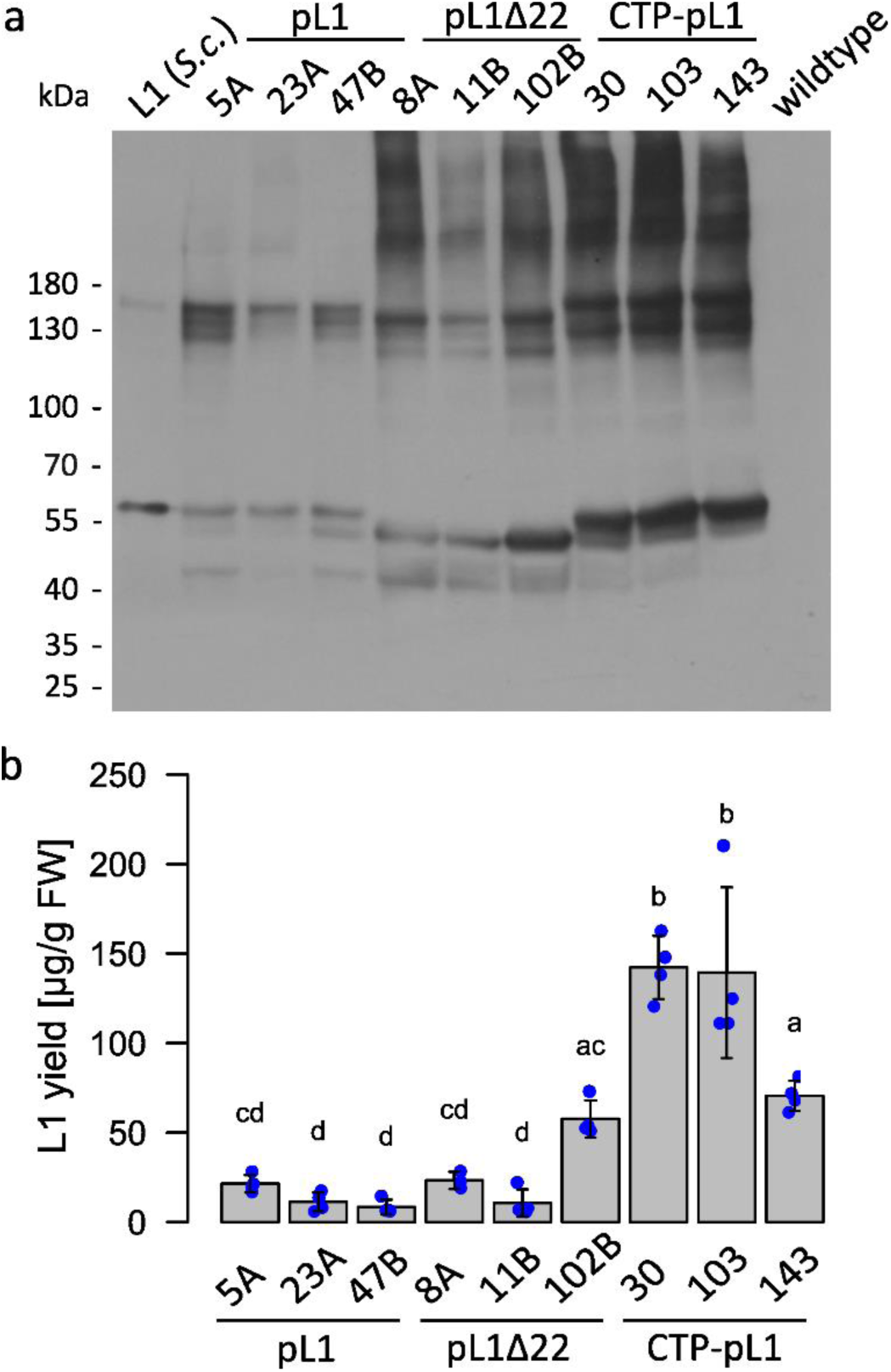
Analysis of HPV-16 L1 production *via* western blot and ELISA. (**a**) For each of the constructs pL1, pL1Δ22, and CTP-pL1 three lines were tested *via* western blot for expression of HPV-16 L1. 50 ng *S. cerevisiae*-produced L1 served as a positive control and Physcomitrella wildtype as a negative control. The proteins for the western blot were separated on a 7.5% TGS polyacrylamide gel under reducing conditions before being transferred to a membrane and developed using CamVir1 antibody (1:5000). (**b**) Production levels of L1 were quantified *via* ELISA. Wildtype background signal was subtracted before analysis *via* one-way ANOVA and post hoc multiple comparisons test (TukeyHSD). Means and standard deviations are calculated based on n ≥ 3 technical replicates from two dilutions and are given per biomass fresh weight (FW). Statistical significance (α = 0.05) is indicated by Compact Letter Display.

Besides the amount of recombinant protein per g FW biomass, the accumulation of biomass is important in biomanufacturing as high product yields per biomass can be trivial if biomass accumulation is slow. To study if recombinant protein amount or subcellular localization of L1 hampers plant growth, we analyzed the time course of biomass accumulation of the nine lines in shaking flasks for 4 weeks.

In our conditions, exponential growth could not be measured. This might be due to the culture conditions, to the rather long intervals between sampling, and/or to the intrinsic growth characteristics of Physcomitrella protonema, which grows by apical cell division (Reski, 1998). Therefore, biomass density was fitted using linear regression analysis model (**Supplementary Figure S6**), using the calculated slope to approximate biomass accumulation rates. Comparison of the slopes between the lines pL1, pL1Δ22 and CTP-pL1 showed pL1 lines accumulated biomass faster than pL1Δ22 (p = <0.0001) and CTP-pL1 (p = 0.0001) lines. If this difference is due to the distinct L1 localization or to lower L1 production in pL1 lines (**Figure 2**) is unclear. Line pL1 47B showed the highest slope of 255 mg dry weight (DW)/L/week, as compared to line CTP-pL1 30 with a slope of 211 mg DW/L/week. However, this difference is negligible considering the yield of pL1 47B is 8 µg L1/g FW as to 142 µg L1/g FW for CTP-pL1 30 (**Figure 2**). Consequently, line CTP- pL1 30 was chosen for VLP production.

### Upscaled L1 production

For production of VLPs, the cultivation of line CTP-pL1 30 was upscaled to 5 L-photobioreactors. The goal here was to identify the best time point for harvesting material used for subsequent L1 purification.

Biomass increased from initially 60 mg DW/L to around 870 mg DW/L at day 4 (**Figure 3**). L1 production remained constant during the first 4 days, in the range of the yield observed for flasks. Upon addition of the phytohormone auxin (NAA) on day 4, L1 levels increased on day 6 and peaked at day 7 at 332 µg/g FW, 130% higher as compared to day 0. Afterwards, L1 production dropped sharply to levels prior to auxin addition. Biomass continued to accumulate until day 11, after which it dropped slightly. These conditions, including day 7 as the ideal harvesting time point, were used for further L1 production.

**Figure 3.**
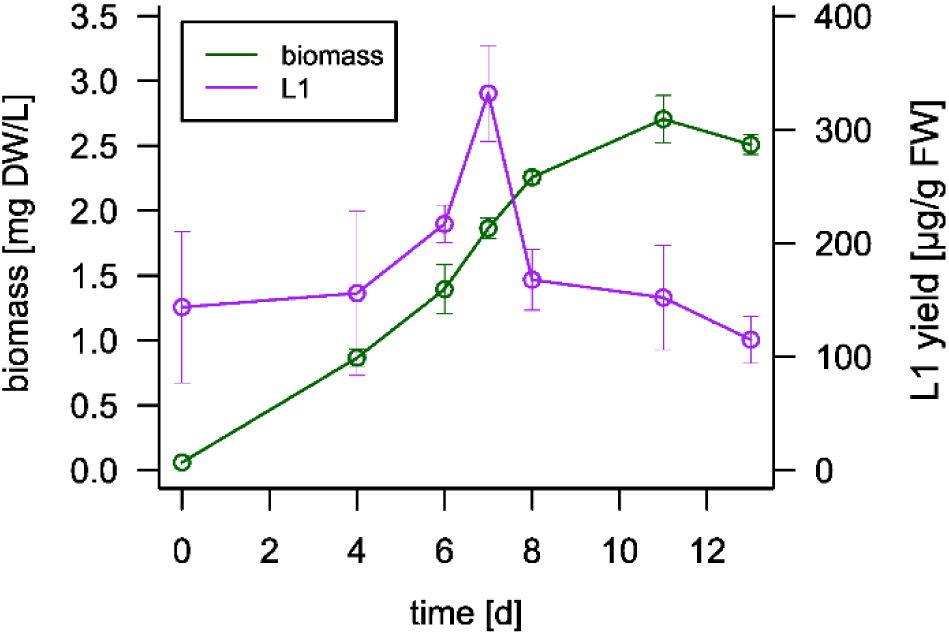
Analysis of biomass accumulation and L1 production of line CTP-pL1 30 in a 5 L- photobioreactor. The bioreactor was inoculated with 60 mg dry weight (DW)/L and monitored over a course of 13 days. On day 4, 5 µM auxin (NAA) was added. L1 production peaked at day 7 at 332 µg/g fresh weight (FW). Biomass and L1 measurements are based on n ≥ 3 technical replicates. Production levels of L1 were quantified *via* ELISA.

**Figure 4.**
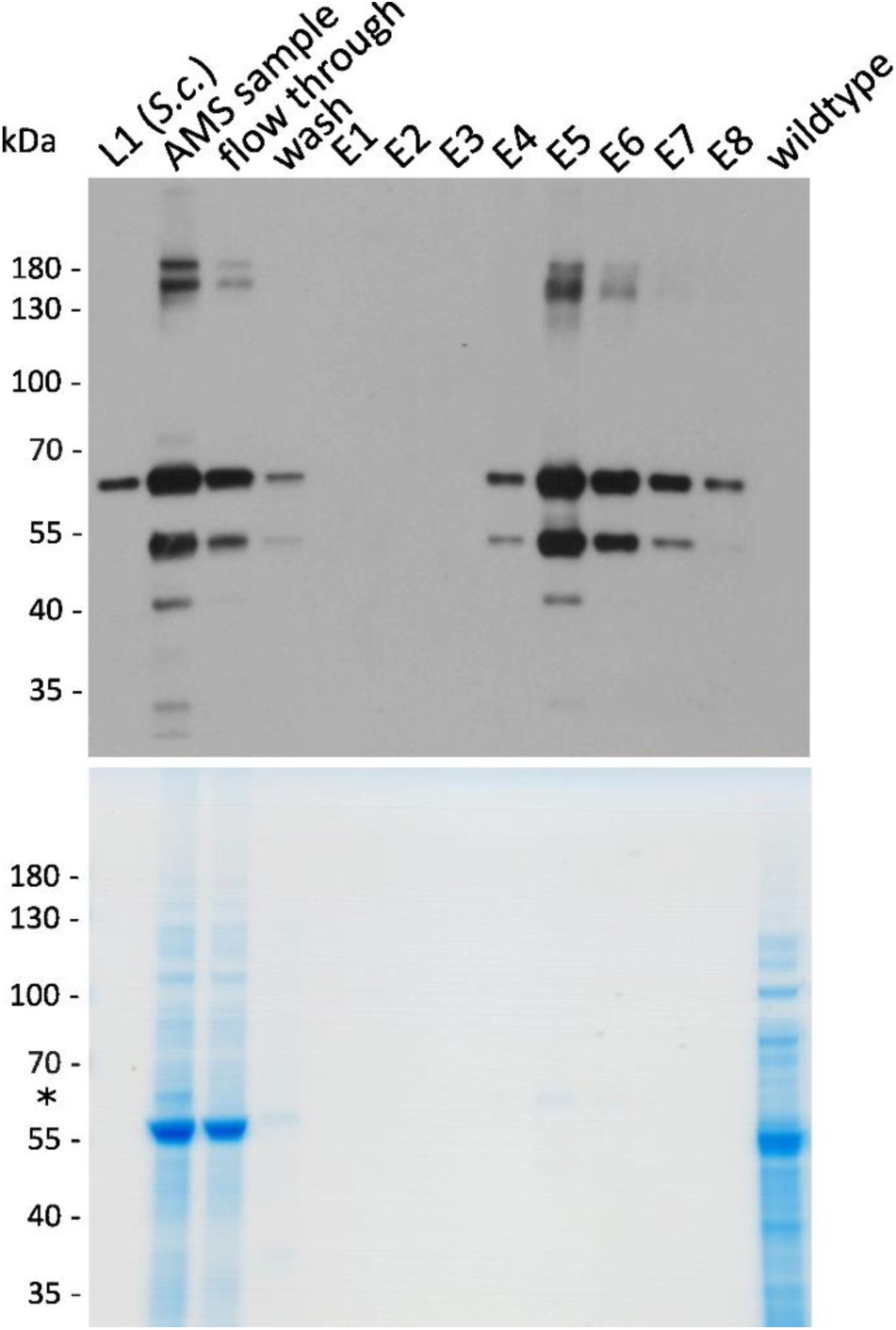
Two-step purification removes bulk of cellular protein. Analysis of L1 and residual cellular host proteins during purification under reducing conditions *via* western blot and Coomassie-stained SDS-PAGE (12% Bis-Tris gel). After precipitation with 45% (w/v) AMS, the filtered sample was subjected to cation exchange chromatography (CaEx) for final purification. Column-bound L1 eluted at 1 M NaCl in fractions E4 – E8. The asterisk in the Coomassie-stained SDS-PAGE marks the band referring to HPV-16 L1. 50 ng *S. cerevisiae*-produced L1 served as a positive control and Physcomitrella wildtype as a negative control.

### Two-step purification removes bulk of cellular proteins

Purification of L1 was performed using ammonium-sulfate (AMS) precipitation, followed by cation exchange (CaEx) chromatography. While AMS removes the bulk of cellular proteins, CaEx removes residual proteins and concentrates the sample.

We tested the efficacy of 40%, 45% and 50% (w/v) AMS to precipitate L1 from an extract containing the soluble protein fraction from CTP-pL1 30 plant material produced in the bioreactor. We found 45% AMS to be sufficient to effectively remove L1 from the supernatant, whereas at 40% AMS small amounts of L1 remained in the supernatant and at 50% AMS higher amounts of cellular protein were precipitated together with L1 (**Supplementary Figure S7**). In preparation for chromatographic purification, the precipitated proteins were resuspended in CaEx binding buffer and dialyzed overnight in binding buffer to remove residual AMS. Cloudiness of samples indicated a reduced solubility of precipitated proteins, and undissolved matter was removed by filtration. The relative amount of L1 and host protein was evaluated *via* western blots and Coomassie-staining, respectively (**Supplementary Figure S7**). Samples after filtration showed a slight reduction in host protein content, but, except for the sample at 40% AMS, no reduction in L1. Overall, precipitation at 45% AMS was selected, which is in line with purifications of yeast-produced L1 (Park et al., 2008; Kim et al., 2010).

Next, CaEx chromatography was used to remove residual cellular proteins. The bulk of cellular proteins did not bind to the column and was detected in the flow through (**Figure 4**, **Supplementary Figure S8**). Notably, a large amount of L1 did not bind to the column and was detected in the flow through. Different binding buffer compositions were tested to increase L1 binding to the column, e.g. lowering the pH from 7.2 to 6.8, lowering the concentration of NaCl from 0.137 M to 0.068 M, addition of 5% (v/v) glycerol, and re-loading of flow-through fraction. Nevertheless, a large part of L1 did not bind to the column and was found in the flow through. During column wash with 5 mL at 0.3 M NaCl, small amounts of contaminating proteins and L1 were washed off the column. For elution, NaCl was increased to 1 M NaCl and 0.5 mL fractions were collected, in total 4 mL. L1 eluted from the column in fractions E4 – E8 (**Figure 4**), which were pooled for further analysis.

### Capsomers and VLPs

Next, we performed a dis- and reassembly protocol on the pooled elution fractions, as it yields more uniform and stable VLPs, which are less prone to aggregation (McCarthy et al., 1998; Mach et al., 2006). First, pooled elution fractions were dialyzed at low salt concentrations, high pH and DTT, that results in the disassembly of VLPs or other multi-pentameric L1-based structures into capsomers. Second, samples were dialyzed at high salt, low pH and in the absence of reducing agent, that leads to the reassembly of capsomers into VLPs.

To assess the conformation of L1 throughout the process, dot blots were performed using different monoclonal antibodies. CamVir1 detects a linear epitope in both denatured and native L1 and served as a control. In contrast, H16.V5 detects only conformational epitopes in both, L1 capsomers and VLPs, but not monomeric or denatured L1.

The dot blot showed the presence of L1 in all CTP-pL1 30-derived samples throughout the purification process, as well as yeast-derived denatured and native L1 (**Figure 5**). In comparison, H16.V5 detects L1 in the native positive control and in all CTP-pL1 30-derived samples, indicating the presence of at least capsomers, if not VLPs, already in the raw extract. Further, conformational epitopes seem to be stable throughout AMS precipitation, CaEx chromatography, disassembly, and reassembly. However, the reassembly sample exhibits a weaker signal for H16.V5 as compared to CamVir1.

**Figure 5.**
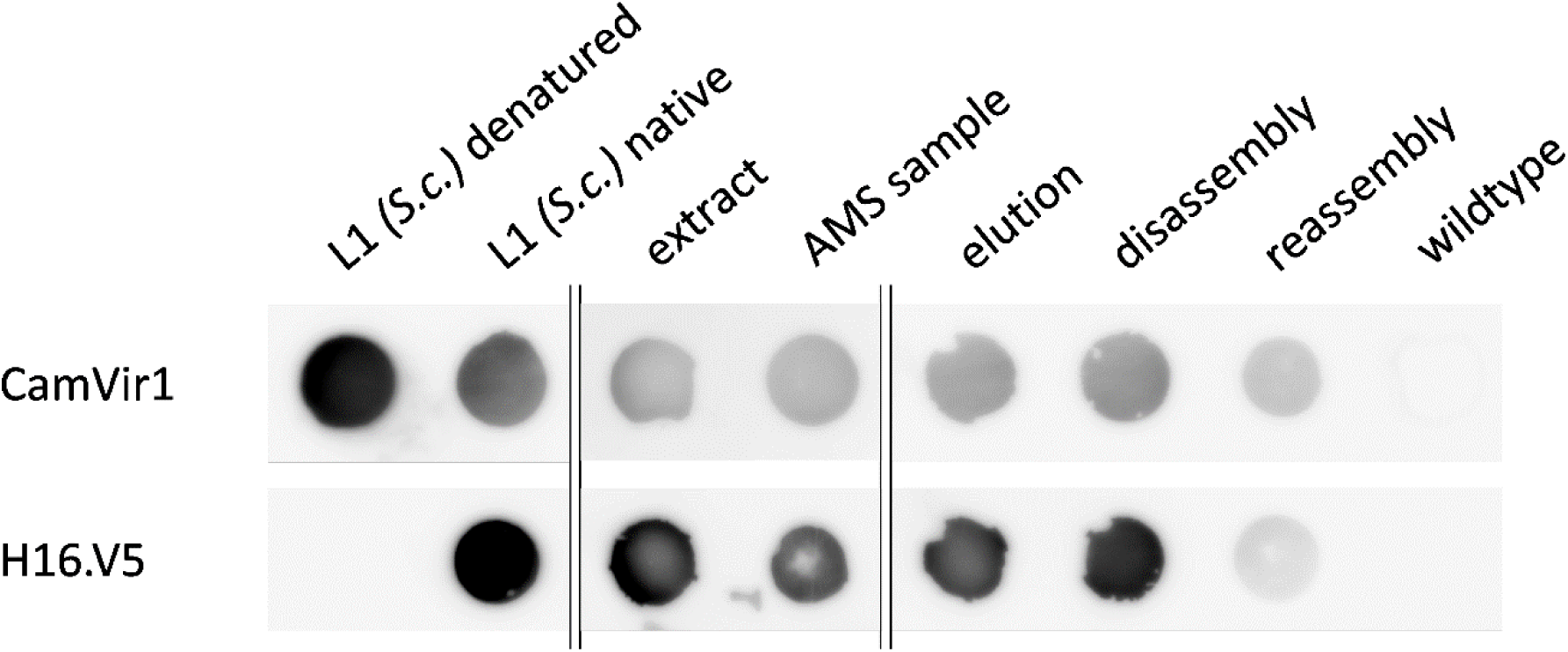
Moss-produced HPV-16 L1 assembles into capsomers, which are stable throughout purification. Dot blots were performed using two different antibodies to analyze the formation of capsomers in yeast-produced L1 and samples from CTP-pL1 30 purification (extract, after AMS precipitation, elution) and dis- and reassembly. The CamVir1 antibody detects a linear epitope in both denatured and native L1. The neutralizing antibody H16.V5 detects specifically conformational epitopes in L1 capsomers and VLPs. Positive H16.V5 signals indicate HPV-16 L1 monomers to assemble at least into capsomers, *in vivo* or after extraction, which are stable throughout the purification. The experiment only allows qualitative but not quantitative comparison since sample concentrations were adjusted individually. ‘L1 denatured’ was incubated at 95 °C for 5 min, all other samples were kept at 4 °C. A wildtype sample was used as a negative control.

The assembly of L1 into VLPs was verified with Transmission Electron Microscopy (TEM). As a control, yeast-produced L1 was analyzed without prior treatment, after disassembly, and after reassembly. In the untreated sample, particles of approx. 60 nm in size were detected. After disassembly, structures of 5 – 10 nm in size were observed, corresponding in size and appearance to capsomers observed for yeast-derived L1 as part of this study (**Figure 6b**) and elsewhere (McCarthy et al., 1996). VLPs reshape upon reassembly treatment, demonstrating the successful dis- and reassembly of yeast-derived L1 VLPs. Notably, reassembled VLPs are heterogenous in size and shape, despite the controlled dis- and reassembly (**Figure 6**).

**Figure 6.**
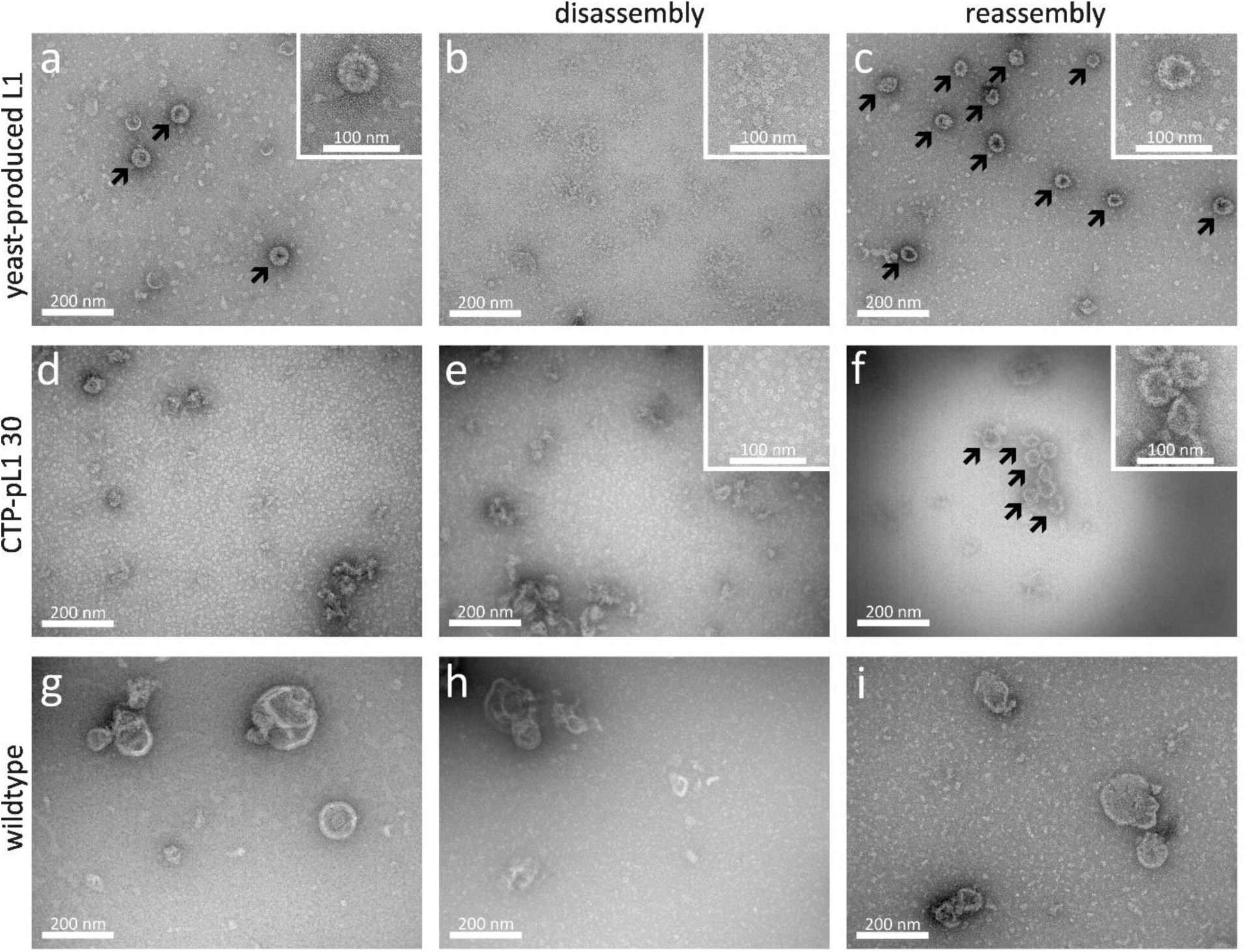
Analysis of dis- and reassembled VLPs from moss and yeast *via* transmission electron microscopy (TEM). Yeast-derived L1 was analyzed untreated (**a**), after disassembly (**b**), and after reassembly (**c**) *via* TEM. CTP-pL130 (**d** – **f**) and wildtype (**g** – **i**) extracts were purified *via* AMS precipitation and CaEx chromatography (**d** & **g**), after disassembly (**e** & **h**), and after reassembly (**f** & **i**). Untreated yeast-derived VLPs (**a**) disintegrate into capsomers upon disassembly (**b**) and reshape into VLPs upon reassembly treatment (**c**). TEM shows the presence of moss-derived HPV-16 L1 capsomers in the elution (**d**) and disassembly sample (**e**) and the assembly into VLPs in the reassembly sample (**f**). Capsomer and VLP-like structures are absent from the wildtype control samples. VLPs are indicated by black arrows.

To analyze L1 assembly into VLPs, samples were taken from the pooled CaEx chromatography elution fractions, after disassembly, and after reassembly.

In the elution sample, no VLPs but capsomeric structures were observed, comparable to those in the disassembled yeast sample, which are not present in the corresponding wildtype control (**Figure 6**). The abundance of capsomers in the elution fractions was surprising given the fact that the elution buffer has a high ionic strength, which favors reassembly. However, before column loading, the sample was dialyzed overnight in binding buffer, which favors disassembly. The exposure to high ionic strength during elution might have been too short to allow VLP assembly. In addition, both binding and elution buffer contained DTT, which induces disassembly (McCarthy et al., 1998; Mach et al., 2006). The disassembled CTP-pL1 30-derived sample depicted capsomers as seen in the elution fraction, and for disassembled yeast-derived L1 (**Figure 6**). **Figure 6f** displays fully assembled, moss-derived HPV-16 L1 VLPs. Again, VLPs were heterogenous in size and shape, as seen for reassembled yeast-derived VLPs. Moss-produced VLPs tended to aggregate, which was not observed for reassembled yeast-derived VLPs, but is common for HPV-16 L1 VLPs (Shi et al., 2005). Aggregation might also explain the relatively low signal for the conformational antibody H16.V5 in the sample of reassembled moss-derived VLPs (**Figure 5**). No VLPs were detected in wildtype samples (**Figure 6**).

## Discussion

The goal of this study was to gain proof-of-concept for the production of virus-like particles (VLPs) in the moss plant Physcomitrella by producing HPV-16 L1 VLPs. For this purpose, we generated a coding sequence (CDS) optimized for expression of HPV-16 L1 in Physcomitrella. We further characterized five chloroplast transit peptides (CTPs) and confirmed the localization of L1 in chloroplasts. In addition, we demonstrated chloroplast localization to yield higher L1 levels than targeting the protein to nuclei or cytoplasm. Cultivation in photobioreactors yielded more than 0.3 mg / g FW, which is in the range of other plant-based expression platforms. We established a purification protocol and demonstrated the successful assembly of moss-produced HPV-16 L1 VLPs.

In tobacco plants (*Nicotiana benthamiana*), various strategies were explored for increasing production yields of HPV-16 L1 VLPs, one of which is optimization of codon usage (Maclean et al., 2007; Hitzeroth et al., 2018). Hitzeroth et al. (2008) found that codon optimization of L1 improves transcription, rather than translation. This is due to RNA motifs in the L1 CDS, which post-transcriptionally regulate the virus life cycle and lead to pre-mRNA degradation (Mori et al., 2006; Zhao and Schwartz, 2008). It is argued that codon optimization removes those motifs, thereby increasing mRNA and protein levels, irrespective of whether the CDS was optimized for expression in humans, tobacco (Hitzeroth et al., 2018) or Physcomitrella (this study). Our Physcomitrella-optimized CDS has a GC content of 62.85% as compared to hL1 with 62.25% (Maclean et al., 2007) and the wildtype sequence with 38.27% (Varsani et al., 2003). Combined with the removal of microRNA binding sites and heterosplice sites we obtained the Physcomitrella-optimized pL1 CDS. We argue that pL1 addresses the issues of L1 intrinsic degradation motifs as well as Physcomitrella-specific posttranscriptional degradation.

Besides codon optimization, research focused on localization signals and transit peptides. The most common strategy is sequestration to chloroplasts, either *via* nuclear transformation and posttranslational import into chloroplasts (Maclean et al., 2007; Matic et al., 2011; Pineo et al., 2013; Zahin et al., 2016; Hitzeroth et al., 2018; Naupu et al., 2020; Muthamilselvan et al., 2023) or *via* transplastomic lines (Fernández-San Millán et al., 2008; Lenzi et al., 2008). Here, we experimentally characterized five Physcomitrella CTPs, of which we used one (CTPc5) for sequestration of L1 to chloroplasts. Employing the HPV-16 native NLS, L1 was targeted to nuclei, whereas the removal of the NLS resulted in cytoplasmic L1. Earlier studies confirmed functionality of the NLS in mammalian cells (Leder et al., 2001; Bissa et al., 2015). However, the NLS failed to localize L1 to nuclei in tobacco (Šmídková et al., 2010). Our observation of the HPV-16 native NLS targeting L1 to nuclei in Physcomitrella is another example for a conserved cross-kingdom functionality of localization signals between moss and mammals (Gitzinger et al., 2009).

Western blots of L1-producing Physcomitrella lines showed a variety of bands, both smaller and larger than the expected 56 kDa or 54 kDa. A comparable banding pattern was observed when expressing L1 in tobacco chloroplasts (Fernández-San Millán et al., 2008). Likewise, yeast-produced L1 exhibited bands with the size of L1 dimers and trimers (Kim et al., 2007; Kim et al., 2010). L1 dimers and trimers are based on inter-capsomeric disulfide bridges which stabilize virions and VLPs (Sapp et al., 1998; Buck et al., 2005; Buck et al., 2013). McCarthy et al. (1998) resolved L1 multimers with DTT. Likewise, Buck et al. (2005) demonstrated absence and presence of L1 dimers and trimers in a comparison of reducing and non-reducing SDS-PAGE and western blots, respectively. An investigation of cysteine 175 and cysteine 428 L1 mutants resulted in the absence of bands at 130 kDa and 180 kDa, thereby linking disulfide bridge-based dimers and trimers to the observed bands (Buck et al., 2005). Surprisingly, reducing agents such as DTT, ascorbic acid or β-mercaptoethanol did not result in the disappearance of these bands in our and other studies (Kim et al., 2007; Fernández-San Millán et al., 2008; Kim et al., 2010). This indicates a tolerance of L1 dimers and trimers to reducing agents or a different origin of multimerization observed at 130 kDa and 180 kDa. Castells-Graells and Lomonossoff (2021) observed multimerization of *Nudaurelia capensis* omega virus α-peptide produced in tobacco, irrespective of DTT, but not when produced in insect cells. These authors hypothesize that cross-linkage of α-peptide occurs during extraction and purification and occurs in plant-based expression systems. Regarding the observed bands below 56 kDa, proteolytic degradation was proposed in other studies observing the same phenomenon (Buck et al., 2005; Fernández-San Millán et al., 2008). In tobacco, a shift from bands above 100 kDa in young leaves to more prominent bands at 45 kDa and 20 kDa in older leaves indicated proteolytic degradation of L1 over time (Fernández-San Millán et al., 2008). In our study, bands at 45 kDa are most prominent in pL1Δ22 lines with cytoplasmic L1 and less prominent for plastidic L1. This might result from organelle-specific proteases (Coates at al., 2022).

In Physcomitrella, combining the full-length L1 with a CTP significantly increases L1 yield as compared to full length L1 or L1 without NLS, which is in line with findings in tobacco (Maclean et al., 2007; Zahin et al., 2016). In our study, L1 yields above 0.3 mg / g FW were achieved in the photobioreactor. In tobacco, yields of 0.1 – 1.2 mg / g FW (Maclean et al., 2007; Pineo et al., 2013; Zahin et al., 2016; Hitzeroth et al., 2018) and up to 3 mg / g FW in transplastomic lines have been reported (Fernández-San Millán et al., 2008). The success of targeting L1 to chloroplasts over localization to other organelles may have several reasons. One is that plastid L1 may be protected from proteases absent in chloroplasts (van Wijk, 2024). Besides, CTPs also act on the transcriptional level. Upon expression in tobacco, chimeric L1/L2 transcripts containing a CTP were more abundant than transcripts lacking a CTP, and consequently yielded more protein (Hitzeroth et al., 2018). So far, it is unclear whether this difference is due to enhanced transcription, increased mRNA half-life or changes in mRNA secondary structure.

For purification of L1 from Physcomitrella extracts, we employed a combination of AMS precipitation and CaEx chromatography. We found 45% AMS to be sufficient to efficiently precipitate L1.

Studies from yeast (Park et al., 2008; Kim et al., 2010) and tobacco (Zahin et al., 2016) also used 45% AMS, whereas sometimes 50% AMS was used (Kim et al., 2018). CaEx chromatography showed that a large part of L1 does not bind to the column. Likewise, Zahin et al. (2016) reported tobacco-produced L1 to bind poorly to CaEx columns. Kim et al. (2010) purified yeast-produced L1 by either CaEx or heparin chromatography. CaEx chromatography showed unbound L1 in the flow through, as observed in our study, whereas no L1 was found in the flow through employing heparin chromatography. Pineo et al. (2013) produced L1/L2 chimaeras in tobacco and experienced loss of L1 and insufficient removal of contaminating proteins using heparin chromatography. Heparin is a highly sulfated form of heparan sulfate, a proteoglycan found on host cell surfaces and a requirement for HPV virion attachment and infection (Knappe et al., 2007). However, a prerequisite for heparin binding is the assembly of L1 to capsomers or VLPs (Rommel et al., 2005), meaning monomers and not properly folded L1 are lost. Overall, chromatography yields for VLPs differ and are influenced by different factors, including the production host, the used materials and buffers, and purification prior to chromatography, making it difficult to compare purification strategies across publications. Here, we achieved to remove the bulk of cellular proteins from moss extracts, but experienced a substantial loss of L1 in the flow through. To address this, further optimization of purification might be achieved by combinations with other methods such as heparin chromatography. Our dot blot analysis confirmed the presence of intact capsomers or VLPs in extracts and after AMS precipitation, suggesting heparin chromatography as an option for the second step in purification.

TEM analysis of purified and reassembled CTP-pL1 samples shows VLPs of approx. 60 nm in size. These nanoparticles assembled from capsomers as a result of the reassembly treatment. The absence of VLPs in the elution fractions does not necessarily indicate that dis- and reassembly treatment is a prerequisite for VLP formation from moss-produced L1, but rather that the buffer conditions did not favor assembly after elution. To the best of our knowledge, this is the first report of VLPs produced in a non-vascular plant and of *in vitro* reassembly of HPV-16 L1 VLPs from capsomers produced in any plant system. The VLPs are heterogenous in shape and size. The same is true for yeast-derived VLPs after reassembly, whereas untreated VLPs were not heterogenous. This is surprising, since controlled dis- and reassembly of VLPs can enhance stability and uniformity (McCarthy et al., 1998; Mach et al., 2006; Shi et al., 2007). Besides high ionic strength and low pH buffer, inter-pentameric disulfide bridges positively influence VLP stability and uniformity (Buck et al., 2005). This could be achieved through prolonged incubation of cleared cell lysates before purification or addition of oxidizing agents, which accelerate disulfide bridge formation.

However, prolonged incubation causes aggregation of HPV-16 L1 VLPs, as observed here, but could be addressed by improved formulation (Shi et al., 2005).

While the moss plant Physcomitrella is an established production host for biopharmaceuticals, we aim to expand its application to other markets such as the cosmetics industry (Munoz et al., 2024) and material sciences at the interface with biomedicine (Milferstaedt et al., 2025; Ramezaniaghdam et al., 2025). In this context, the present study serves as a proof-of-concept for the production of a relatively simple but medicinally relevant VLP in Physcomitrella. Future efforts may focus on more complex eVLPs that require proper N-glycosylation or complex folding and assembly; a specialty niche well suited for the moss-based expression platform.

## Acknowledgements

We thank the Life Imaging Centre (LIC) of the University of Freiburg for support with confocal microscopy. We are grateful for technical assistance by Agnes Novakovic, Britta Rothgänger, Christine Glockner and Richard Haas and to Anne Katrin Prowse for language editing.

## Author contributions

PAN: Conceptualization, Formal analysis, Funding acquisition, Investigation, Methodology, Visualization, Writing – original draft, Writing – review & editing

MCW: Investigation, Methodology, Writing – review & editing SC: Investigation, Methodology, Writing – review & editing

CET: Investigation, Methodology, Formal analysis, Writing – review & editing MRF: Investigation, Formal analysis, Writing – review & editing

SNWH: Investigation, Formal analysis, Writing – review & editing JP: Writing – original draft, Writing – review & editing

ELD: Writing – original draft, Writing – review & editing

HTS: Conceptualization, Resources, Writing – review & editing

RR: Supervision, Resources, Writing – original draft, Writing – review & editing

## Data availability statement

Nucleotide sequence data was submitted to GenBank under accession numbers PV029542 (pL1) and PV068547(CTPc5). Moss lines and vectors were submitted to the International Moss Stock Centre (IMSC) under accession numbers 40963 (pL1 5A), 40964 (pL1 23A), 40965 (pL1 47B), 40966 (pL1Δ22 8A), 40967 (pL1Δ22 11B), 40968 (pL1Δ22 102B), 40969 (CTP-pL1 30), 40970 (CTP-pL1 103), 40971 (CTP-pL1 143), 1840 (pAct5_linker_eGFP_35ST_hpt), 2038 (pAct5_CTP_linker_eGFP_35ST_hpt), 1193 (pAct5_Citrine_nosT_hpt), 2249 (pAct5_pL1-Citrine_nosT_hpt), 2250 (pAct5_pL1Δ22- Citrine_nosT_hpt), 2251 (pAct5_CTP-pL1-Citrine_nosT_hpt), 1997 (pAct5_pL1_35ST_hpt), 1998 (pAct5_pL1Δ22_35ST_hpt) and 2048 (pAct5_CTP-pL1_35ST_hpt).

## Short legends for Supporting Information

Supplementary Figure S1 **Vector maps**

Supplementary Figure S2 **Generation of Physcomitrella-optimized (p)L1**

Supplementary Figure S3 **Characterization of CTPs by analysis of subcellular eGFP localization**

Supplementary Figure S4 **Analysis of CDS integrity in L1-producing lines**

Supplementary Figure S5 **Analysis of the effect of reducing agents on L1 band patterning**

Supplementary Figure S6 **Comparison of biomass accumulation rates** Supplementary Figure S7 **Ammonium sulfate precipitation** Supplementary Figure S8 **Chromatogram of Cation Exchange purification** Supplementary Table S1 **List of primers**

Supplementary Table S2. Predicted chloroplast transit peptides for six selected proteins in Physcomitrella

---

**Supplementary Figure S1.**
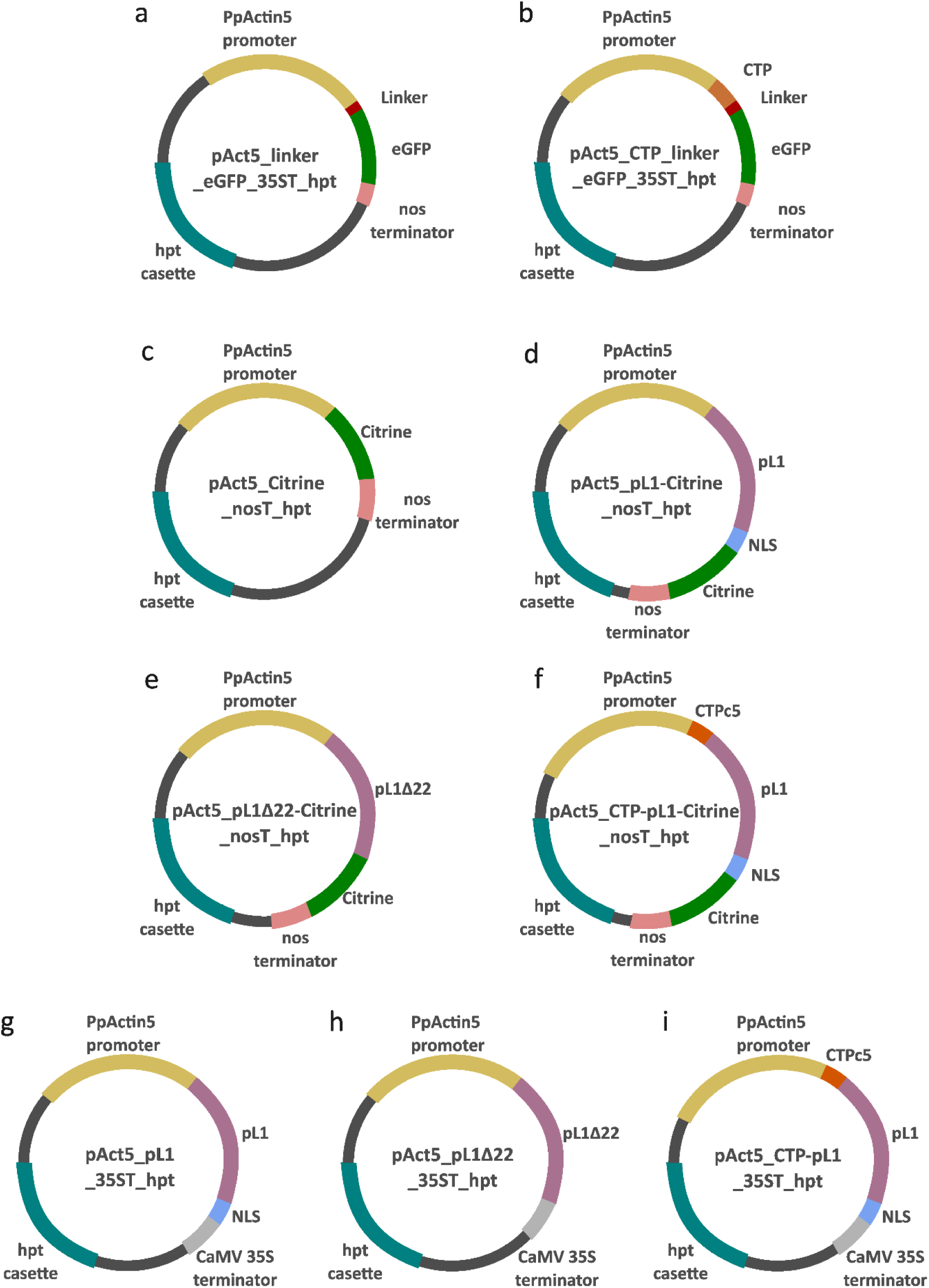
– Vector maps **List of all vectors used in this study.** Vectors **a** & **b** were used for the characterization of chloroplast transit peptides (CTPs). Vector **a** served as a control and was used as a cloning template for inserting different CTP candidates. Vectors **c**, **d**, **e** & **f** were used for the analysis of L1 subcellular targeting. Vector **c** served as a control and was used as a cloning template for inserting either pL1, pL1Δ22 or CTP-pL1 CDS (**d**, **e** & **f**, respectively). Vectors **g**, **h** & **i** were used for generation of stable transgenic Physcomitrella lines. The vectors contain either the full length pL1 including the native nuclear localization signal (NLS) (**g**), a truncated version lacking the NLS (**h**), or the full-length version including the native NLS and an N-terminal chloroplast transit peptide (**i**). All vectors contain the PpActin5 promoter (yellow), the CaMV 35S terminator (grey), and a hygromycin (hpt)(turquoise) resistance cassette for selection. Other elements are the GGGGGA linker (red), CTP (orange), NLS (blue) and eGFP or Citrine (green). Genetic elements are not depicted true to scale. Vector sizes: 7260 bp (**a**), 7461 bp (**b**), 7299 bp (**c**), 8802 bp (**d**), 8736 bp (**e**), 9021 bp (**f**), 7998 bp (**g**), 7932 bp (**h**), 8217 bp (**i**).

**Supplementary Figure S2.**
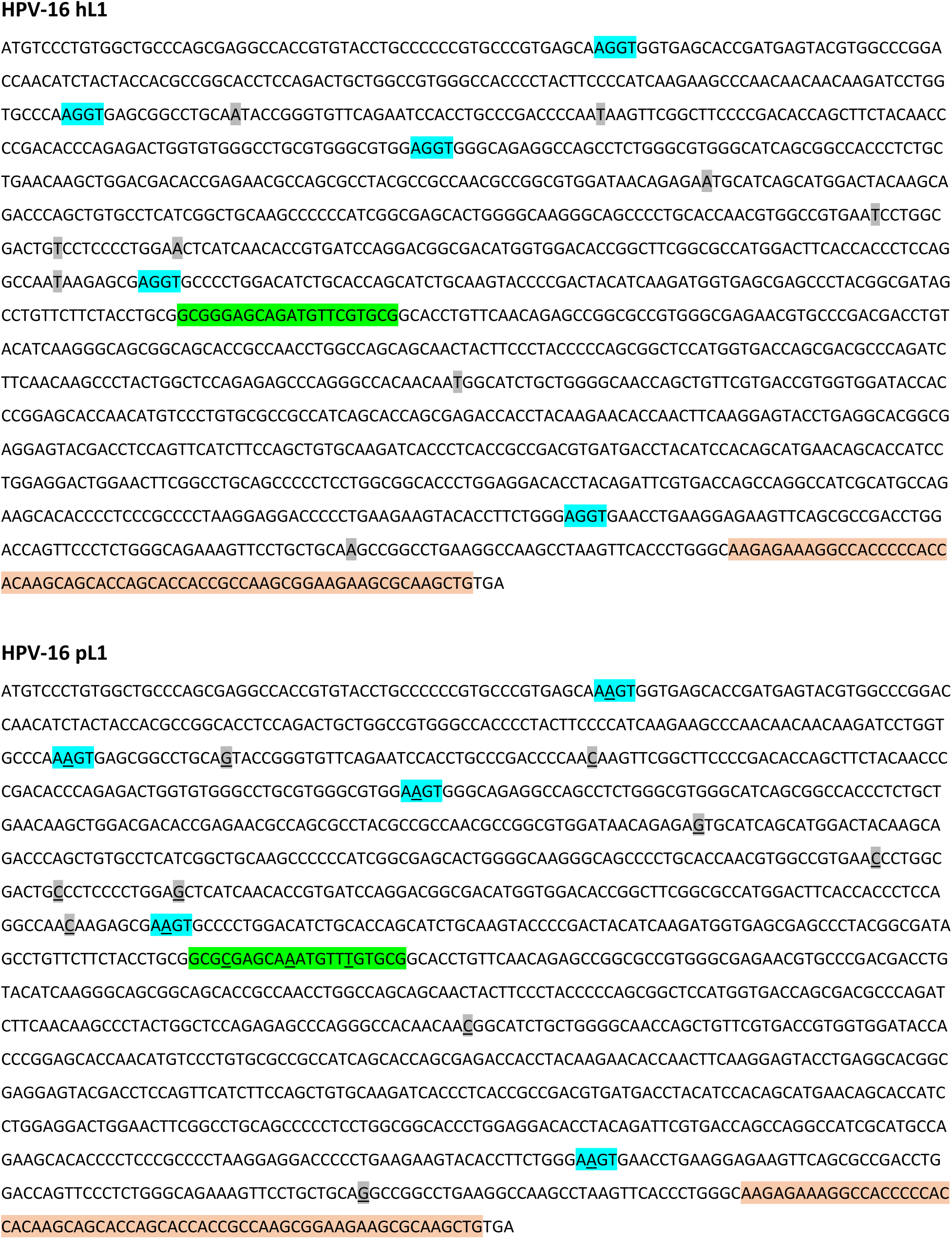
- Generation of Physcomitrella-optimized (p)L1 HPV-16 hL1 ATGTCCCTGTGGCTGCCCAGCGAGGCCACCGTGTACCTGCCCCCCGTGCCCGTGAGCAAGGTGGTGAGCACCGATGAGTACGTGGCCCGGA CCAACATCTACTACCACGCCGGCACCTCCAGACTGCTGGCCGTGGGCCACCCCTACTTCCCCATCAAGAAGCCCAACAACAACAAGATCCTGGTGCCCAAGGTGAGCGGCCTGCAATACCGGGTGTTCAGAATCCACCTGCCCGACCCCAATAAGTTCGGCTTCCCCGACACCAGCTTCTACAACCCCGACACCCAGAGACTGGTGTGGGCCTGCGTGGGCGTGGAGGTGGGCAGAGGCCAGCCTCTGGGCGTGGGCATCAGCGGCCACCCTCTGCTGAACAAGCTGGACGACACCGAGAACGCCAGCGCCTACGCCGCCAACGCCGGCGTGGATAACAGAGAATGCATCAGCATGGACTACAAGCAGACCCAGCTGTGCCTCATCGGCTGCAAGCCCCCCATCGGCGAGCACTGGGGCAAGGGCAGCCCCTGCACCAACGTGGCCGTGAATCCTGGCGACTGTCCTCCCCTGGAACTCATCAACACCGTGATCCAGGACGGCGACATGGTGGACACCGGCTTCGGCGCCATGGACTTCACCACCCTCCAGGCCAATAAGAGCGAGGTGCCCCTGGACATCTGCACCAGCATCTGCAAGTACCCCGACTACATCAAGATGGTGAGCGAGCCCTACGGCGATAGCCTGTTCTTCTACCTGCGGCGGGAGCAGATGTTCGTGCGGCACCTGTTCAACAGAGCCGGCGCCGTGGGCGAGAACGTGCCCGACGACCTGTACATCAAGGGCAGCGGCAGCACCGCCAACCTGGCCAGCAGCAACTACTTCCCTACCCCCAGCGGCTCCATGGTGACCAGCGACGCCCAGATCTTCAACAAGCCCTACTGGCTCCAGAGAGCCCAGGGCCACAACAATGGCATCTGCTGGGGCAACCAGCTGTTCGTGACCGTGGTGGATACCACCCGGAGCACCAACATGTCCCTGTGCGCCGCCATCAGCACCAGCGAGACCACCTACAAGAACACCAACTTCAAGGAGTACCTGAGGCACGGCGAGGAGTACGACCTCCAGTTCATCTTCCAGCTGTGCAAGATCACCCTCACCGCCGACGTGATGACCTACATCCACAGCATGAACAGCACCATCCTGGAGGACTGGAACTTCGGCCTGCAGCCCCCTCCTGGCGGCACCCTGGAGGACACCTACAGATTCGTGACCAGCCAGGCCATCGCATGCCAGAAGCACACCCCTCCCGCCCCTAAGGAGGACCCCCTGAAGAAGTACACCTTCTGGGAGGTGAACCTGAAGGAGAAGTTCAGCGCCGACCTGGACCAGTTCCCTCTGGGCAGAAAGTTCCTGCTGCAAGCCGGCCTGAAGGCCAAGCCTAAGTTCACCCTGGGCAAGAGAAAGGCCACCCCCACCACAAGCAGCACCAGCACCACCGCCAAGCGGAAGAAGCGCAAGCTGTGA **HPV-16 pL1** ATGTCCCTGTGGCTGCCCAGCGAGGCCACCGTGTACCTGCCCCCCGTGCCCGTGAGCAA<u>A</u>GTGGTGAGCACCGATGAGTACGTGGCCCGGAC CAACATCTACTACCACGCCGGCACCTCCAGACTGCTGGCCGTGGGCCACCCCTACTTCCCCATCAAGAAGCCCAACAACAACAAGATCCTGGT GCCCAA<u>A</u>GTGAGCGGCCTGCA<u>G</u>TACCGGGTGTTCAGAATCCACCTGCCCGACCCCAA<u>C</u>AAGTTCGGCTTCCCCGACACCAGCTTCTACAACCC CGACACCCAGAGACTGGTGTGGGCCTGCGTGGGCGTGGA<u>A</u>GTGGGCAGAGGCCAGCCTCTGGGCGTGGGCATCAGCGGCCACCCTCTGCT GAACAAGCTGGACGACACCGAGAACGCCAGCGCCTACGCCGCCAACGCCGGCGTGGATAACAGAGA<u>G</u>TGCATCAGCATGGACTACAAGCA GACCCAGCTGTGCCTCATCGGCTGCAAGCCCCCCATCGGCGAGCACTGGGGCAAGGGCAGCCCCTGCACCAACGTGGCCGTGAA<u>C</u>CCTGGC GACTG<u>C</u>CCTCCCCTGGA<u>G</u>CTCATCAACACCGTGATCCAGGACGGCGACATGGTGGACACCGGCTTCGGCGCCATGGACTTCACCACCCTCCA GGCCAA<u>C</u>AAGAGCGA<u>A</u>GTGCCCCTGGACATCTGCACCAGCATCTGCAAGTACCCCGACTACATCAAGATGGTGAGCGAGCCCTACGGCGATA GCCTGTTCTTCTACCTGCGGCG<u>C</u>GAGCA<u>A</u>ATGTT<u>T</u>GTGCGGCACCTGTTCAACAGAGCCGGCGCCGTGGGCGAGAACGTGCCCGACGACCTG TACATCAAGGGCAGCGGCAGCACCGCCAACCTGGCCAGCAGCAACTACTTCCCTACCCCCAGCGGCTCCATGGTGACCAGCGACGCCCAGAT CTTCAACAAGCCCTACTGGCTCCAGAGAGCCCAGGGCCACAACAA<u>C</u>GGCATCTGCTGGGGCAACCAGCTGTTCGTGACCGTGGTGGATACCA CCCGGAGCACCAACATGTCCCTGTGCGCCGCCATCAGCACCAGCGAGACCACCTACAAGAACACCAACTTCAAGGAGTACCTGAGGCACGGCGAGGAGTACGACCTCCAGTTCATCTTCCAGCTGTGCAAGATCACCCTCACCGCCGACGTGATGACCTACATCCACAGCATGAACAGCACCATCCTGGAGGACTGGAACTTCGGCCTGCAGCCCCCTCCTGGCGGCACCCTGGAGGACACCTACAGATTCGTGACCAGCCAGGCCATCGCATGCCA GAAGCACACCCCTCCCGCCCCTAAGGAGGACCCCCTGAAGAAGTACACCTTCTGGGA<u>A</u>GTGAACCTGAAGGAGAAGTTCAGCGCCGACCTG GACCAGTTCCCTCTGGGCAGAAAGTTCCTGCTGCA<u>G</u>GCCGGCCTGAAGGCCAAGCCTAAGTTCACCCTGGGCAAGAGAAAGGCCACCCCCAC CACAAGCAGCACCAGCACCACCGCCAAGCGGAAGAAGCGCAAGCTGTGA **Human codon-optimized HPV-16 (h)L1 CDS and Physcomitrella-optimized HPV-16 (p)L1 CDS.** Grey marked nucleotides were changed in HPV-16 pL1 through codon usage optimization by the tool physCO (Top et al., 2021). Turquoise marked motifs refer to the Physcomitrella alternative splicing recognition sites. The green marked segment refers to a predicted miRNA binding site identified by the tool psRNATarget (Dai and Zhao, 2011; Dai et al., 2018). Light orange colored segments represent the nuclear localization signal. Nucleotide changes are underlined in the HPV-16 pL1 sequence. (hL1, GenBank accession number DQ067889; Maclean et al., 2007).

**Supplementary Figure S3.**
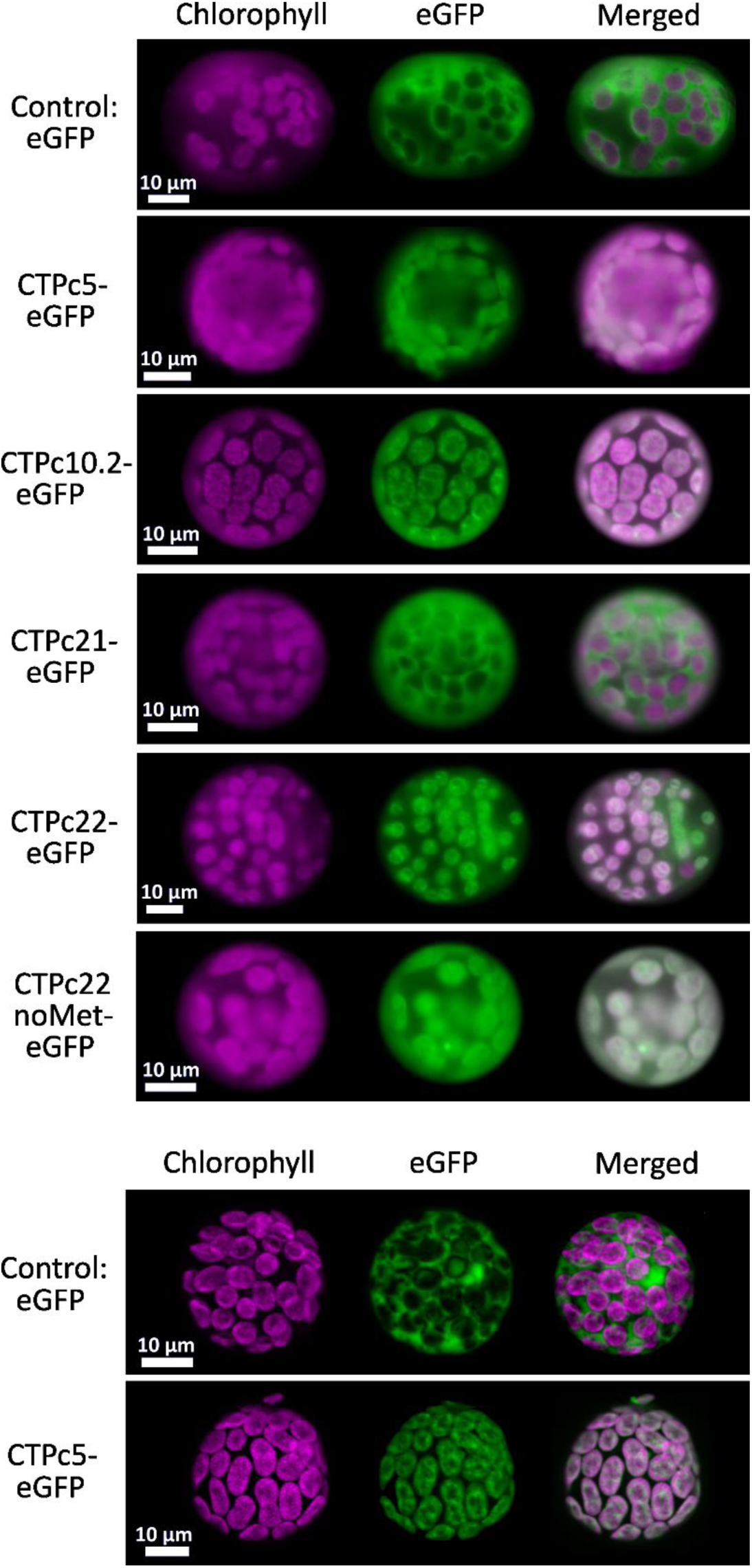
- Characterization of CTPs by analysis of subcellular eGFP localization **Microscopy pictures of Physcomitrella protoplasts transformed with CTP-eGFP constructs.** Samples were analyzed using a rhodamine and a FITC filter to identify chloroplast (magenta) and eGFP localization, respectively, as well as a merge of the two signals. For sample CTPc21-eGFP no eGFP signal could be observed in chloroplasts. For samples CTPc5-eGFP, CTPc10-eGFP, CTPc22-eGFP and CTPc22noMet-eGFP green fluorescence signal is observed in chloroplasts. eGFP lacking a CTP localized to the cytoplasm and served as a negative control. The experiment was repeated two times. CTPc5 was examined a third time using confocal microscopy. Scale bars represent 10 µm.

**Supplementary Figure S4.**
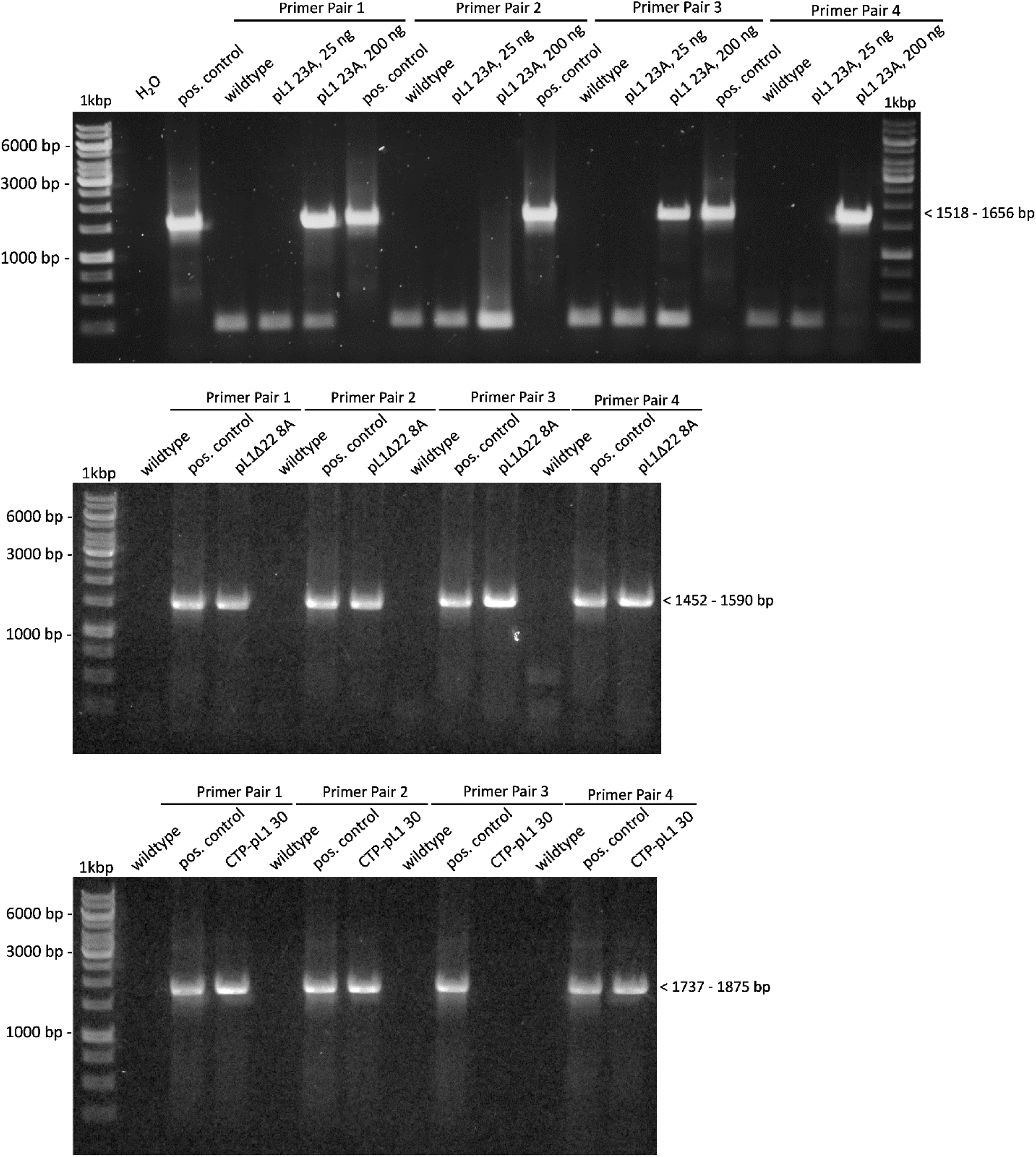
– Analysis of CDS integrity in L1-producing lines **Analysis of rearrangements in the pL1 CDS using RT PCR.** Lines pL1 23A, pL1Δ22 8A and CTP-pL1 30 were checked for DNA or RNA rearrangements by performing PCR on cDNA using four different primer pairs. Primer pairs were selected to amplify the entire cDNA. The forward primer binds at the start codon and is the same in each primer pair (pL1_fwd_1 for lines pL1 23A & pL1Δ22 8A, CTP_pL1_fwd_1 for line CTP- pL1 30). The reverse primers are different in each pair and bind at the 3’ end of the CDS or within the terminator (**Supplementary Table S1**). For line pL1 23A usage of 25 ng or 200 ng cDNA in PCR reactions was compared. Following experiments used 200 ng. Absence of distinct bands of a size smaller than the expected size is indicative for no larger rearrangements on both DNA and RNA level. No amplicon is visible for lines pL1 23A using Primer Pair 2 and CTP-pL1 30 using Primer Pair 3. The constructs used for transformation of Physcomitrella protoplasts (**Supplementary Figure S1 g - i**) served as a positive control.

**Supplementary Figure S5.**
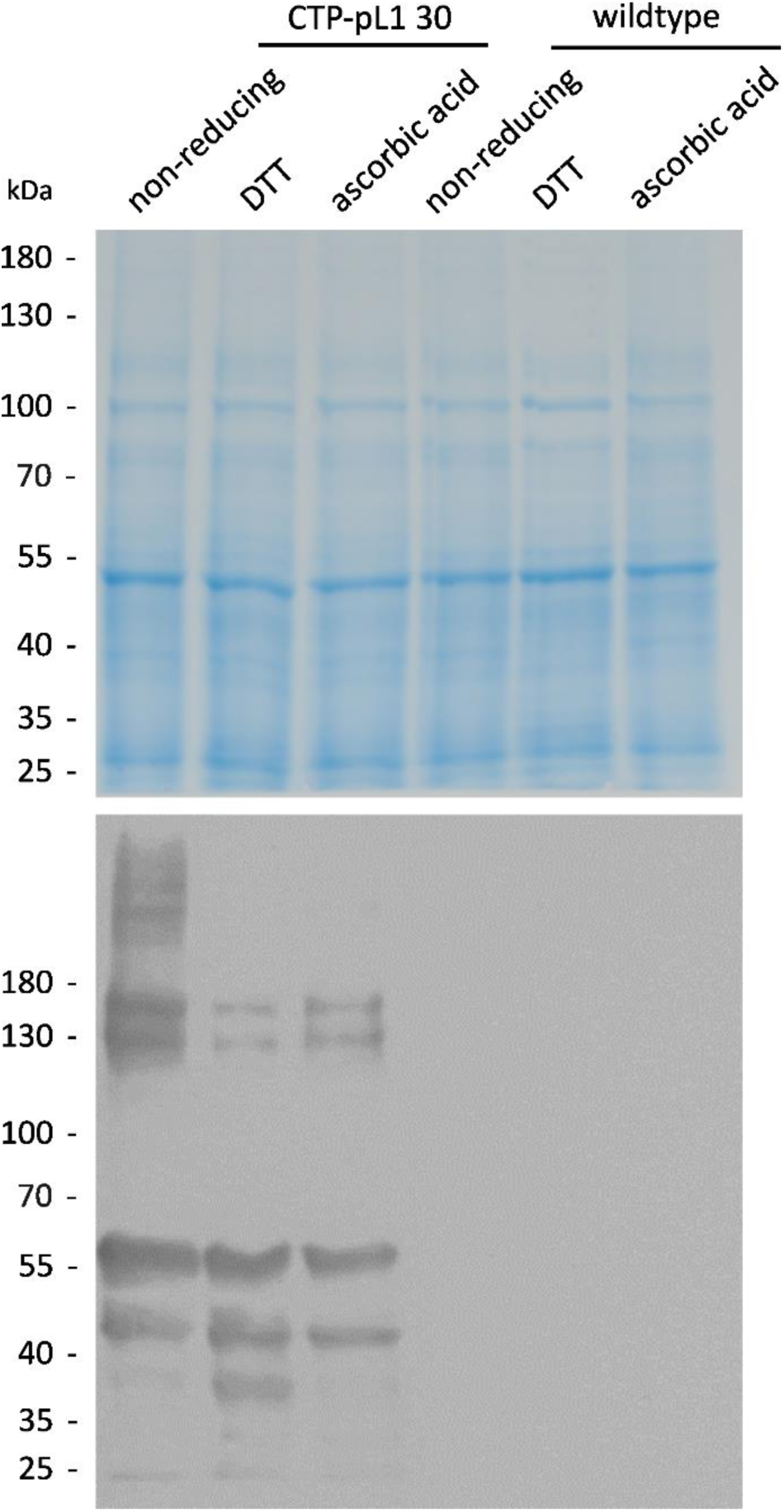
– Analysis of the effect of reducing agents on L1 band patterning **Analysis of the effect of reducing agents on L1 band patterning in western blots.** Samples of line CTP- pL1 30 and wildtype were extracted without reducing agent, with 50 mM DTT or 40 mM ascorbic acid. The proteins for the western blot were separated on a 7.5% TGS polyacrylamide gel under reducing conditions (5 mM DTT) before being transferred to a membrane and developed using CamVir1 antibody (1:5000).

**Supplementary Figure S6.**
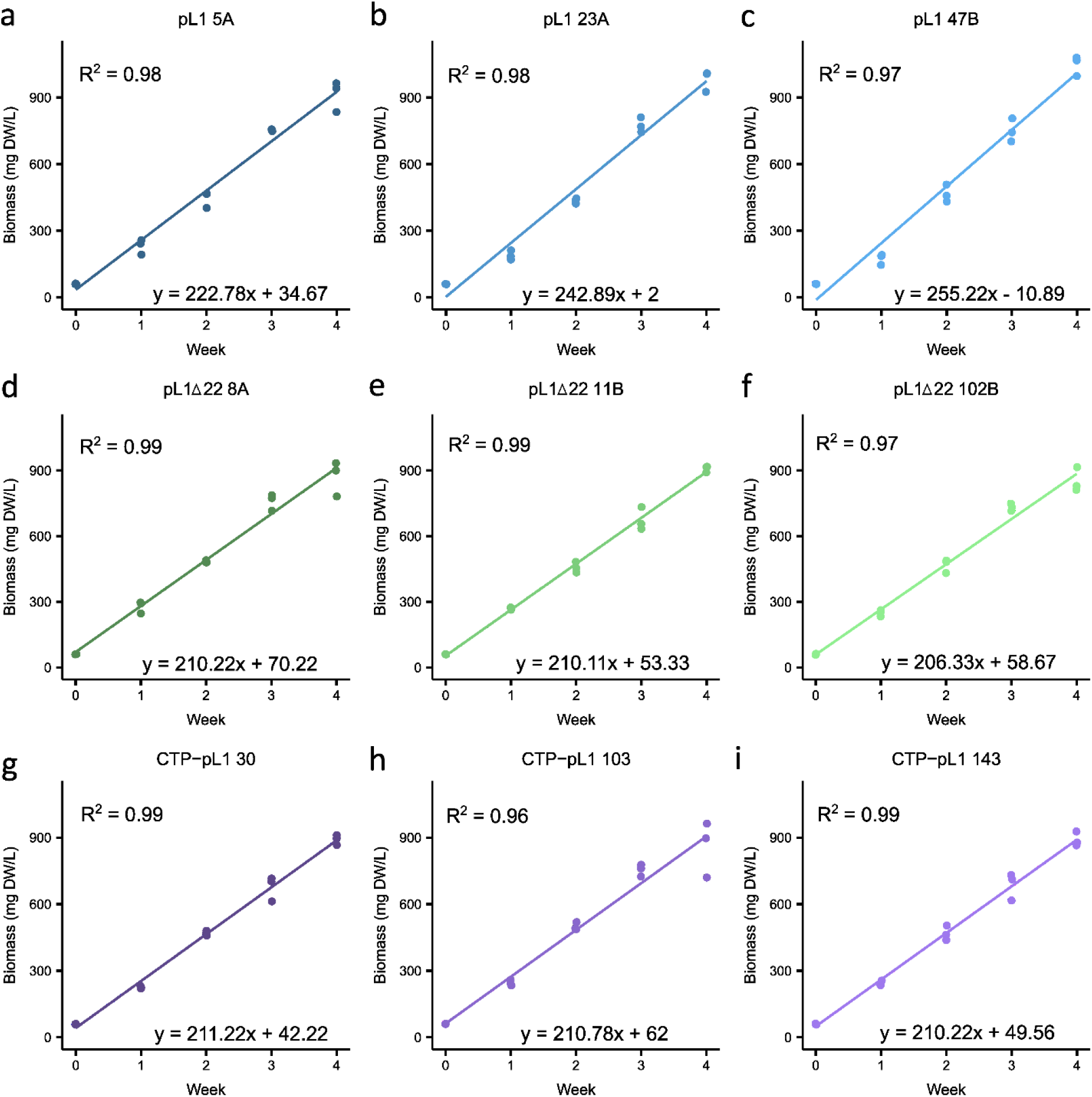
– Comparison of biomass accumulation rates **Analysis of biomass accumulation rates among nine Physcomitrella lines of three different constructs.** Biomass accumulation rates were observed over a course of 4 weeks for lines pL1 5A, pL1 23A, pL1 47B (blue); pL1Δ22 8A, pL1Δ22 11B, pL1Δ22 102B (green) and CTP-pL1 30, CTP-pL1 103, CTP-pL1 143 (purple). The data was plotted using linear regression analysis and all slopes were analysed using ANOVA and post hoc multiple comparisons test (TukeyHSD). Comparison of the slopes between the constructs of pL1, pL1Δ22 and CTP-pL1 showed pL1 lines accumulated biomass faster than pL1Δ22 (p = <0.0001) and CTP- pL1 (p = 0.0001) lines. Per week n = 3 biological samples were taken.

**Supplementary Figure S7.**
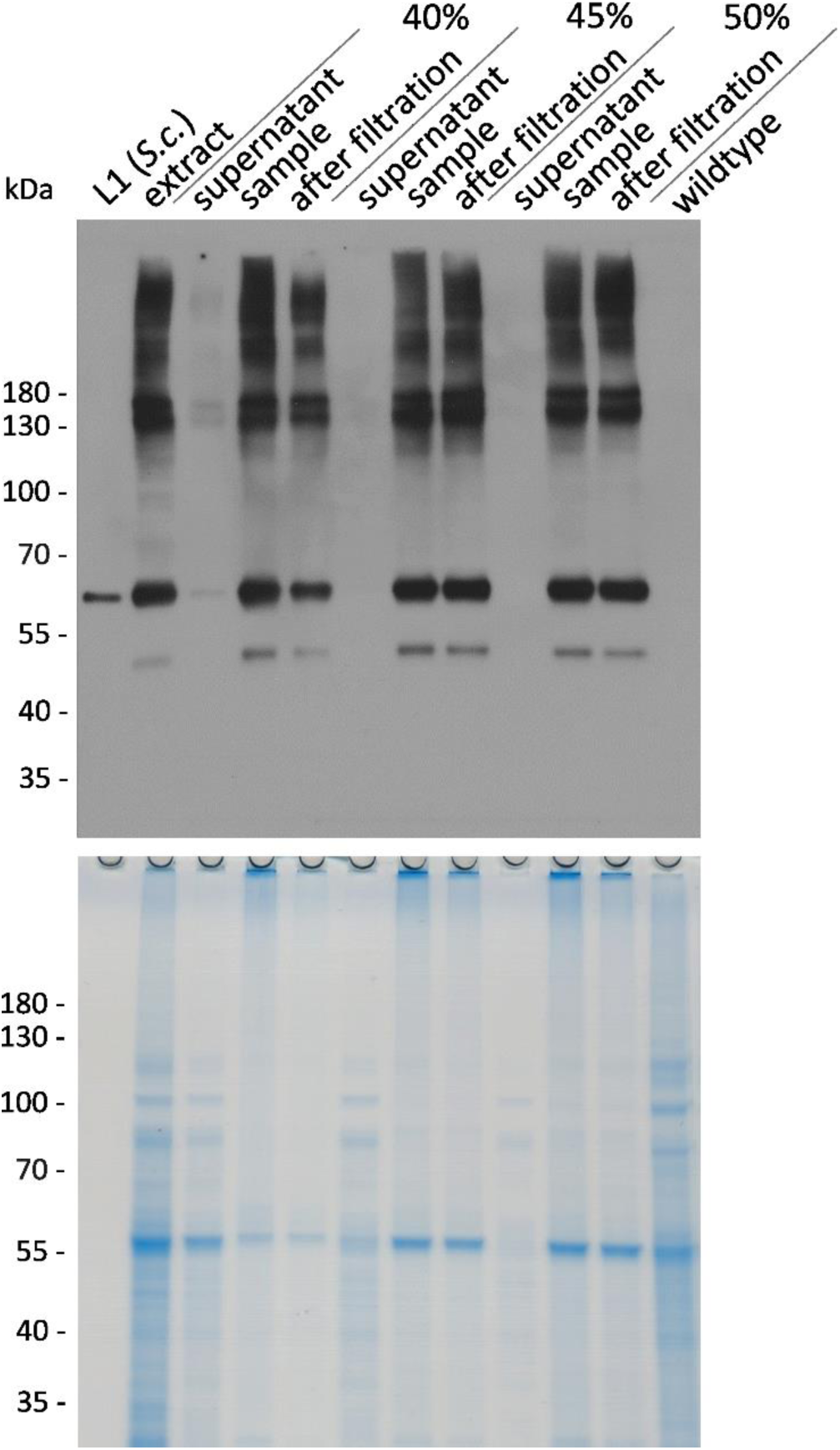
– Ammonium sulfate precipitation **Ammonium sulfate precipitation of L1 from moss extracts.** Samples from the supernatant and the resuspended and dialyzed pellet before and after filtration were analyzed for the presence of L1 content and residual host proteins under reducing conditions *via* western blot and Coomassie-stained gel, respectively. CTP-pL1 30 extracts were precipitated with either 40%, 45%, or 50% (w/v) ammonium sulfate (AMS). After centrifugation, the supernatant and the precipitate were taken up and dialyzed against cation exchange chromatography binding buffer to remove excess AMS. Non-dissolved protein was cleared from the sample using 0.22 µm filters. 45% AMS was found to be sufficient for full precipitation of L1 from crude extracts. 30 ng *S. cerevisiae*-produced L1 served as a positive control and Physcomitrella wildtype as a negative control.

**Supplementary Figure S8.**
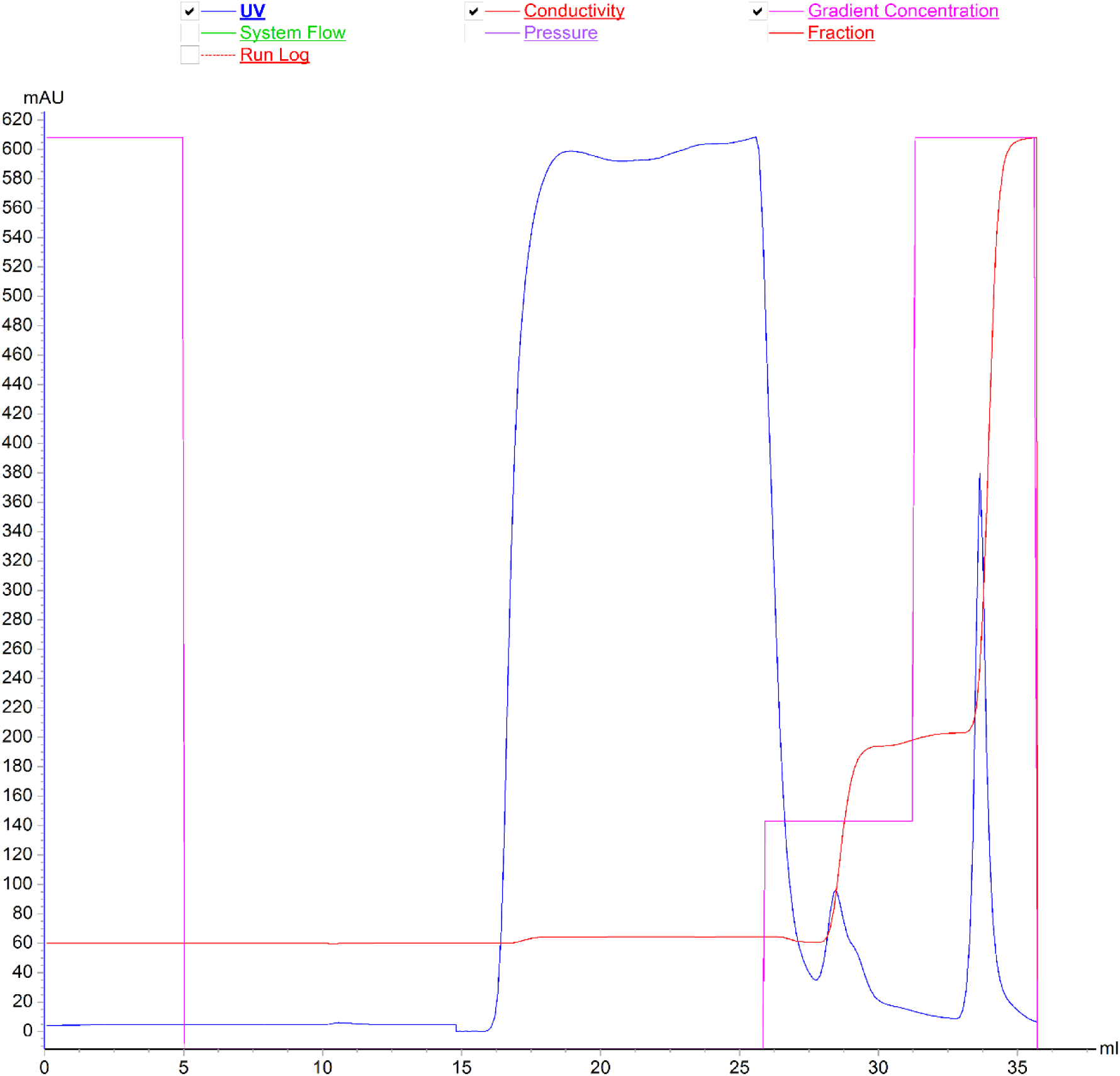
– Chromatogram of Cation Exchange purification **Chromatogram of HPV-16 L1 purification from moss using cation exchange chromatography.** The chromatogram shows the UV absorbance, indicative for protein concentration, in blue, gradient concentration in pink, and conductivity, indicative for salt concentration, in red. The y axis depicts milli- Absorbance Units and the x axis depicts the volume in mL. The large peak between 16 – 26 mL shows the sample application or flow through, the following small peak at 28 mL the wash fraction and the larger peak at 33 – 35 mL the elution fractions.

**Supplementary Table S1.**
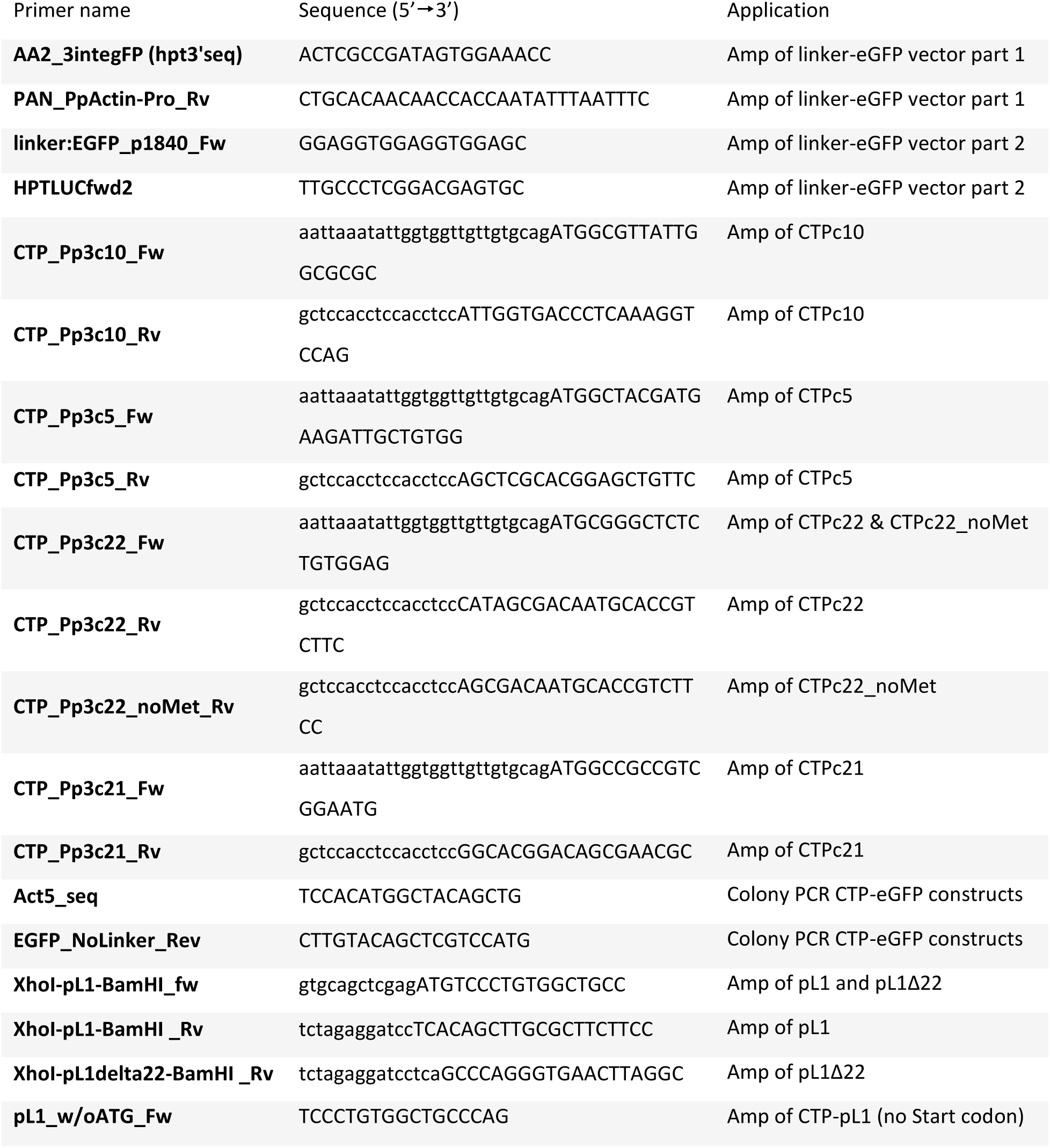
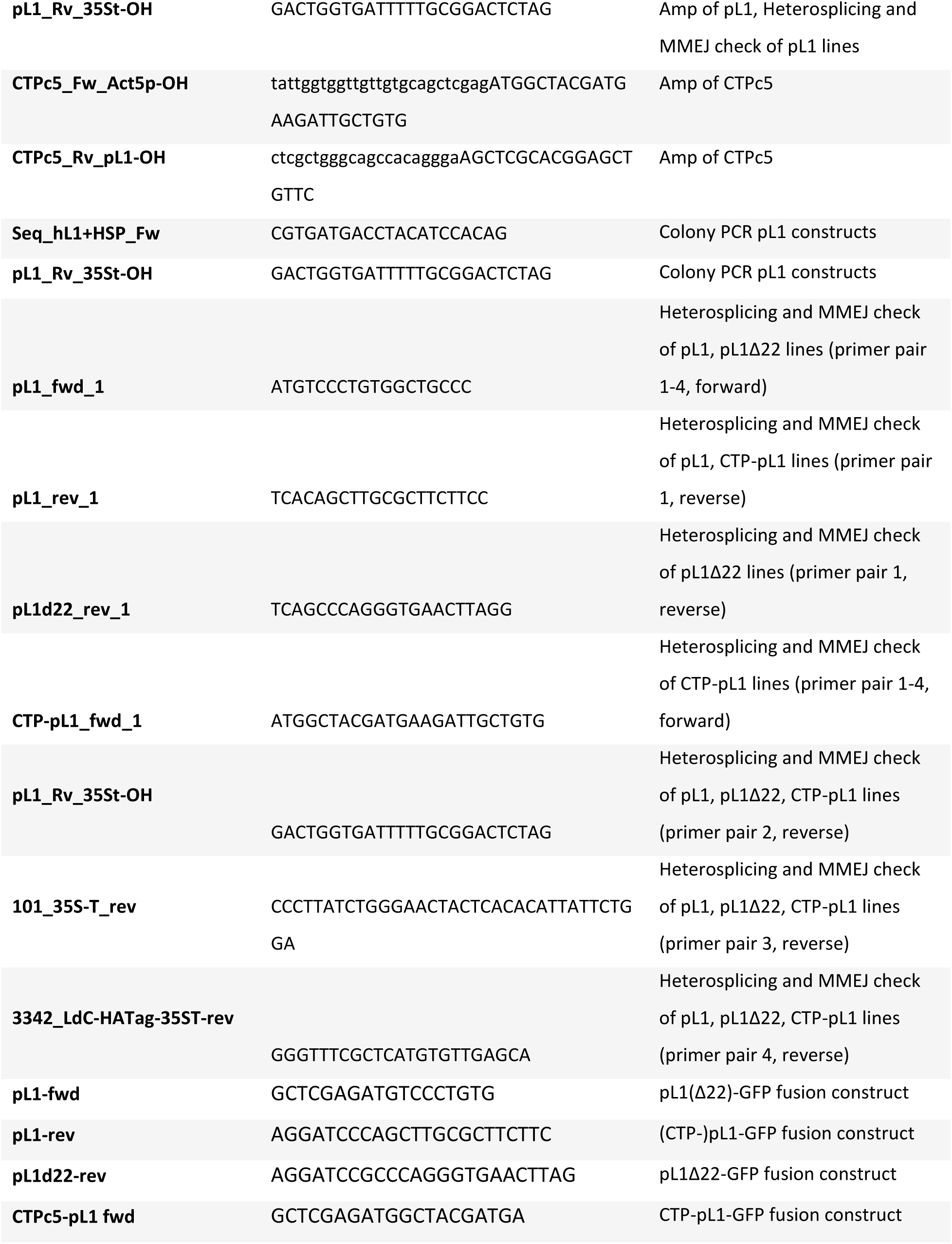
– List of primers Capital letters refer to the part of the primer binding to the template whereas lower case letters represent overhangs added to the template via PCR. Abbreviations: fwd or fw (forward), rev or rv (reverse), CTP (chloroplast transit peptide), MMEJ (micro homology-mediated end joining), GFP (green fluorescent protein), OH (overhang (lower case letters)).

**Supplementary Table S2.**
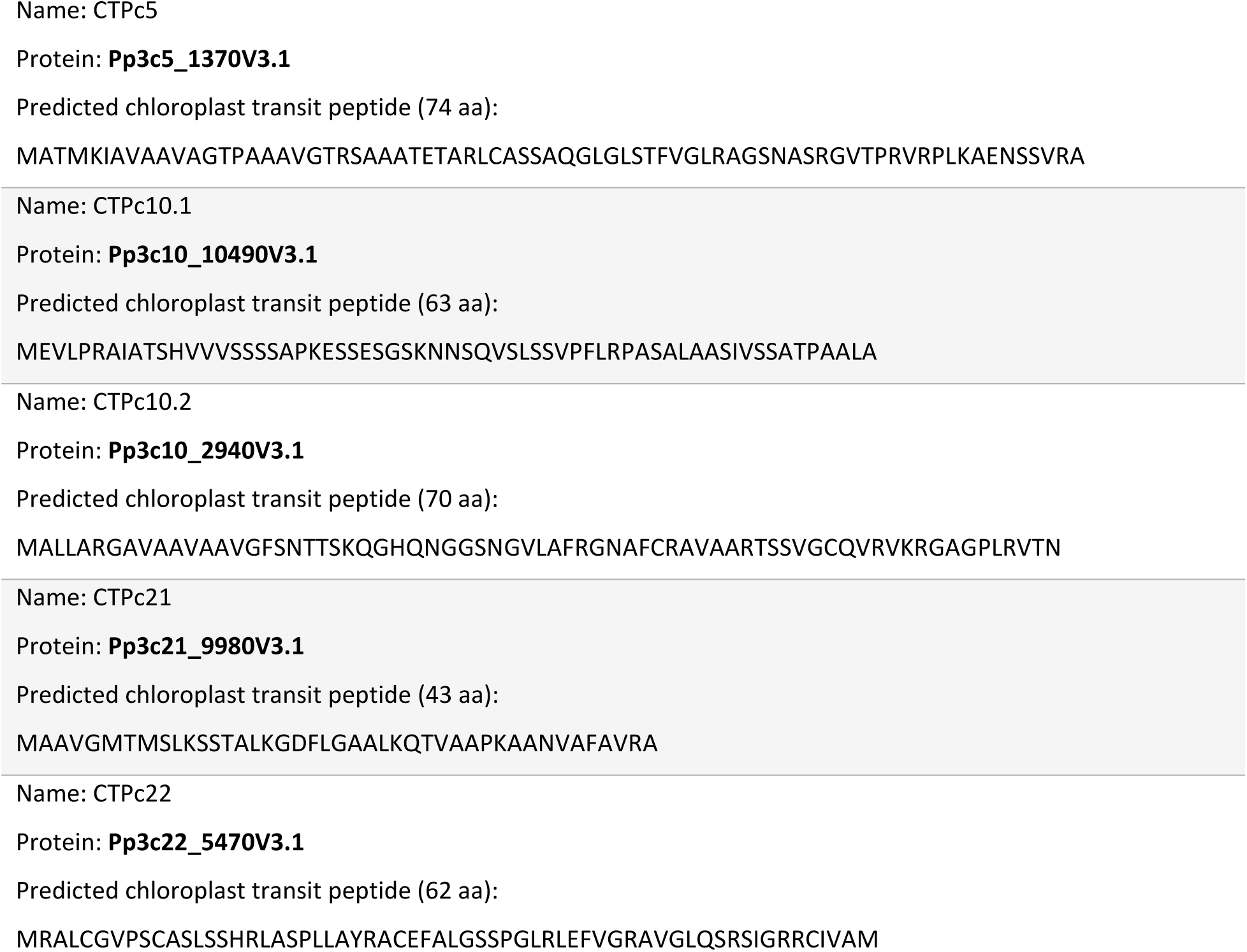
– Predicted chloroplast transit peptides for six selected proteins in Physcomitrella. The predictions were performed with TargetP-2.0; Armenteros et al. (2019).

## References

1. Abel WO, Knebel W, Koop HU, Marienfeld JR, Quader H, Reski R, Schnepf E, Sporlein B. A cytokinin-sensitive mutant of the moss, *Physcomitrella patens*, defective in chloroplast division. Protoplasma (1989) 152:1–13. doi: 10.1007/BF01354234

2. Adams A, Hendrikse M, Rybicki EP, Hitzeroth II. Optimal size of DNA encapsidated by plant produced human papillomavirus pseudovirions. Virology (2023) 580:88–97. doi: 10.1016/j.virol.2023.02.003

3. Armenteros JJA, Salvatore M, Emanuelsson O, Wither O, von Heijne G, Elofsson A, Nielsen H. Detecting sequence signals in targeting peptides using deep learning. Life Sci All (2019) 2:e201900429. doi: 10.26508/lsa.201900429

4. Baker TS, Newcomb WW, Olson NH, Cowsert LM, Olson C, Brown JC. Structures of bovine and human papillomaviruses. Analysis by cryoelectron microscopy and three-dimensional image reconstruction. Biophys J (1991) 60:1445–1456. doi: 10.1016/S0006-3495(91)82181-6

5. Benvenuto E, Broer I, D’Aoust MA, Hitzeroth I, Hundleby P, Menassa R, Oksman-Caldentey KM, Peyret H, Salgueiro S, Saxena P, Stander J, Warzecha H, Ma J. Plant molecular farming in the wake of the closure of Medicago Inc. Nature Biotech (2023) 41:893–894. doi: 10.1038/s41587-023-01812-w

6. Biemelt S, Sonnewald U, Galmbacher P, Willmitzer L, Müller M. Production of human papillomavirus type 16 virus-like particles in transgenic plants. J Virol (2003) 77:9211–9220. doi: 10.1128/JVI.77.17.9211-9220.2003

7. Bissa M, Zanotto C, Pacchioni S, Volonté L, Venuti A, Lembo D, De Giuli Morghen C, Radaelli A. The L1 protein of human papilloma virus 16 expressed by a fowlpox virus recombinant can assemble into virus-like particles in mammalian cell lines but elicits a non-neutralising humoral response. Antiviral Res (2015) 116:67–75. doi: 10.1016/j.antiviral.2015.01.012

8. Bohlender LL, Parsons J, Hoernstein SNW, Rempfer C, Ruiz-Molina N, Lorenz T, Rodríguez Jahnke F, Figl R, Fode B, Altmann F, Reski R, Decker EL. Stable protein sialylation in Physcomitrella. Front Plant Sci (2020) 11:610032. doi: 10.3389/FPLS.2020.610032

9. Bohlender LL, Parsons J, Hoernstein SNW, Bangert N, Rodriguez-Jahnke F, Reski R, Decker EL. Unexpected arabinosylation after humanization of plant protein N-glycosylation. Front Bioeng Biotech (2022) 10:838365. 10.3389/fbioe.2022.838365

10. Buck CB, Thompson CD, Pang YYS, Lowy DR, Schiller JT. Maturation of papillomavirus capsids. J Virol (2005) 79:2839–2846. doi: 10.1128/JVI.79.5.2839-2846.2005

11. Buck CB, Day PM, Trus BL. The papillomavirus major capsid protein L1. Virol (2013) 445:169–174. doi: 10.1016/j.virol.2013.05.038

12. Burns CC, Diop OM, Sutter RW, Kew OM. Vaccine-derived polioviruses. J Infect Dis (2014) 210:S283-S293. doi: 10.1093/INFDIS/JIU295

13. Büttner-Mainik A, Parsons J, Jérôme H, Hartmann A, Lamer S, Schaaf A, Schlosser A, Zipfel PF, Reski R, Decker EL. Production of biologically active recombinant human factor H in Physcomitrella. Plant Biotech J (2011) 9:373–383. doi: 10.1111/J.1467-7652.2010.00552.X

14. Castells-Graells R, Lomonossoff GP. Plant-based production can result in covalent cross-linking of proteins. Plant Biotech J (2021) 19:1095–1097. doi: 10.1111/pbi.13598

15. Cerqueira C, Schiller JT. Papillomavirus assembly: An overview and perspectives. Virus Res (2017) 231:103–107. doi: 10.1016/j.virusres.2016.11.010

16. Chi H, Pepper M, Thomas PG. Principles and therapeutics of adaptive immunity. Cell (2024) 187:2052–2078. doi: doi.org/10.1016/j.cell.2024.03.037

17. Coates RJ, Young MT, Scofield S. Optimising expression and extraction of recombinant proteins in plants. Front Plant Sci (2022) 13:1074531. doi: 10.3389/fpls.2022.1074531

18. Cubas R, Zhang S, Li M, Chen C, Yao Q. Chimeric Trop2 virus-like particles: A potential immunotherapeutic approach against pancreatic cancer. J Immunoth (2011) 34:251–263. doi: CJI.0B013E318209EE72

19. Dai X, Zhao PX. psRNATarget: a plant small RNA target analysis server. Nucl Acids Res (2011) 39:W155–W159. doi: 10.1093/nar/gkr319

20. Dai X, Zhuang, Z, Zhao, PX. psRNATarget: a plant small RNA target analysis server (2017 release). Nucl Acids Res (2018) 46:W49–W54. doi: 10.1093/NAR/GKY316

21. de Martel C, Georges D, Bray F, Ferlay J, Clifford GM. Global burden of cancer attributable to infections in 2018: a worldwide incidence analysis. Lancet Glob Health (2020) 8:e180–e190. doi: 10.1016/S2214-109X(19)30488-7

22. Decker EL, Reski R. Mosses in biotechnology. Curr Op Biotech (2020) 61:21–27. doi: 10.1016/j.copbio.2019.09.021

23. Decker EL, Wiedemann G, Reski R. Gene targeting for precision glyco-engineering: Production of biopharmaceuticals devoid of plant-typical glycosylation in moss bioreactors. Meth Mol Biol (2015) 1321:213–224. doi: 10.1007/978-1-4939-2760-9_15

24. Decker EL, Parsons J, Reski R. Glyco-engineering for biopharmaceutical production in moss bioreactors. Front Plant Sci (2014) 5:346. doi: 10.3389/fpls.2014.00346

25. Eleva (2025). Eleva and 3PBIOVAN sign strategic GMP manufacturing and technology alliance to boost capacity for Eleva’s platform and pipeline. https://elevabiologics.com/eleva-and-3pbiovian-sign-strategic-alliance [Accessed May 21, 2025].

26. Fernández-San Millán A, Ortigosa SM, Hervás-Stubbs S, Corral-Martínez P, Seguí-Simarro JM, Gaétan J, Coursaget P, Veramendi J. Human papillomavirus L1 protein expressed in tobacco chloroplasts self- assembles into virus-like particles that are highly immunogenic. Plant Biotech J (2008) 6:427–441. doi: 10.1111/j.1467-7652.2008.00338.x

27. Gibson DG, Young L, Chuang RY, Venter JC, Hutchison CA, Smith HO. Enzymatic assembly of DNA molecules up to several hundred kilobases. Nature Met (2009) 6:343–345. doi: 10.1038/nmeth.1318

28. Gitzinger M, Parsons J, Reski R, Fussenegger M. Functional cross-kingdom conservation of mammalian and moss (*Physcomitrella patens*) transcription, translation and secretion machineries. Plant Biotech J (2009) 7:73–86. doi: 10.1111/J.1467-7652.2008.00376.X

29. Gremillon L, Kiessling J, Hause B, Decker EL, Reski R, Sarnighausen E. Filamentous temperature-sensitive Z (FtsZ) isoforms specifically interact in the chloroplasts and in the cytosol of *Physcomitrella patens*. New Phyt (2007) 176:299–310. doi: 10.1111/j.1469-8137.2007.02169.x

30. Gupta R, Arora K, Roy SS, Joseph A, Rastogi R, Arora NM, Kundu PK. Platforms, advances, and technical challenges in virus-like particles-based vaccines. Front Immun (2023) 14:1123805. doi: 10.3389/FIMMU.2023.1123805

31. Hasan SS, Sevvana M, Kuhn RJ, Rossmann MG. Structural biology of Zika virus and other flaviviruses. Nat Struc Mol Biol (2018) 25:13–20. doi: 10.1038/s41594-017-0010-8

32. Hector M, Behnke V, Dabrowska-Schlepp, Busch A, Schaaf A, Langmann T, Wolf A. Moss-derived human complement factor H modulates retinal immune response and attenuates retinal degeneration. J Neuroinflam (2025) 22:104. doi: 10.1186/s12974-025-03418-2

33. Hennermann JB, Arash-Kaps L, Fekete G, Schaaf A, Busch A, Frischmuth T. Pharmacokinetics, pharmacodynamics, and safety of moss-aGalactosidase A in patients with Fabry disease. J Inher Metab Dis (2019) 42:527–533. doi: 10.1002/JIMD.12052

34. Hitzeroth II, Chabeda A, Whitehead MP, Graf M, Rybicki EP. Optimizing a human papillomavirus type 16 L1-based chimaeric gene for expression in plants. Front Bioeng Biotech (2018) 6:101. doi: 10.3389/fbioe.2018.00101

35. Hoernstein SNW, Oezdemir B, van Gessel N, Miniera AA, Rogalla von Bieberstein B, Nilges L, Schweikert Farinha J, Komoll R, Glauz S, Weckerle T, Scherzinger F, Rodriguez-Franco M, Mueller-Schuessele S, Reski R. A deeply conserved protease, acylamino acid-releasing enzyme (AARE), acts in ageing in Physcomitrella and Arabidopsis. Comm Biol (2023) 6:61. doi: 10.1038/s42003-023-04428-7

36. Hoernstein SNW, Schlosser A, Fiedler K, van Gessel N, Igloi GL, Lang D, Reski R. A snapshot of the Physcomitrella N-terminome reveals N-terminal methylation of organellar proteins. Plant Cell Rep (2024) 43:250. doi: 10.1007/s00299-024-03329-1

37. Hohe A, Reski R. Optimisation of a bioreactor culture of the moss *Physcomitrella patens* for mass production of protoplasts. Plant Sci (2002) 163: 69–74. doi: 10.1016/S0168-9452(02)00059-6

38. Hohe A, Decker EL, Gorr G, Schween G, Reski R. Tight control of growth and cell differentiation in photoautotrophically growing moss (*Physcomitrella patens*) bioreactor cultures. Plant Cell Rep (2002) 20:1135–1140. doi: 10.1007/s00299-002-0463-y

39. Hohe A, Egener T, Lucht JM, Holtorf H, Reinhard C, Schween G, Reski R. An improved and highly standardized transformation procedure allows efficient production of single and multiple targeted gene- knockouts in a moss, *Physcomitrella patens*. Curr Genet (2004) 44:339–347. doi: 10.1007/s00294-003-0458-4

40. Jones-Rhoades MW, Bartel DP, Bartel B. MicroRNAs and their regulatory roles in plants. Ann Rev Plant Biol (2006) 57:19–53. doi: 10.1146/annurev.arplant.57.032905.105218

41. Kerr H, Herbert AP, Makou E, Abramczyk D, Malik TH, Lomax-Browne H, Yang Y, Pappworth IY, Denton H, Richards A, Marchbank KJ, Pickering MC, Barlow PN. Murine factor H co-produced in yeast with protein disulfide isomerase ameliorated C3 dysregulation in factor H-deficient mice. Front Immun (2021) 12:681098. 10.3389/FIMMU.2021.681098

42. Khraiwesh B, Arif MA, Seumel GI, Ossowski S, Weigel D, Reski R, Frank W. Transcriptional control of gene expression by microRNAs. Cell (2010) 140:111–122. doi: 10.1016/j.cell.2009.12.023

43. Kim SN, Jeong HS, Park SN, Kim HJ. Purification and immunogenicity study of human papillomavirus type 16 L1 protein in *Saccharomyces cerevisiae*. J Virol Met (2007) 139:24–30. doi: 10.1016/j.jviromet.2006.09.004

44. Kim HJ, Kim SY, Lim SJ, Kim JY, Lee SJ, Kim HJ. One-step chromatographic purification of human papillomavirus type 16 L1 protein from *Saccharomyces cerevisiae*. Prot Expr Pur (2010) 70:68–74. doi: 10.1016/j.pep.2009.08.005

45. Kim HS, Jeon JH, Lee KJ, Ko K. N-glycosylation modification of plant-derived virus-like particles: An application in vaccines. BioMed Res Int (2014) 2014:249519. doi: 10.1155/2014/249519

46. Kim HJ, Cho SY, Park MH, Kim HJ. Comparison of the size distributions and immunogenicity of human papillomavirus type 16 L1 virus-like particles produced in insect and yeast cells. Arch Pharm Res (2018) 41:544–553. doi: 10.1007/s12272-018-1024-4

47. Kirnbauer R, Booy F, Cheng N, Lowy DR, Schiller JT. Papillomavirus L1 major capsid protein self-assembles into virus-like particles that are highly immunogenic. Proc Nat Acad Sci USA (1992) 89:12180–84. doi: 10.1073/pnas.89.24.12180

48. Knappe M, Bodevin S, Selinka HC, Spillmann D, Streeck RE, Chen XS, Lindahl U, Sapp M. Surface-exposed amino acid residues of HPV16 L1 protein mediating interaction with cell surface heparan sulfate. J Biol Chem (2007) 282:27913–22. 10.1074/jbc.M705127200

49. Koprivova A, Stemmer C, Altmann F, Hoffmann A, Kopriva S, Gorr G, Reski R, Decker EL. Targeted knockouts of *Physcomitrella* lacking plant-specific immunogenic N-glycans. Plant Biotech J (2004) 2:517–523. doi: 10.1111/J.1467-7652.2004.00100.X

50. Lamprecht RL, Kennedy P, Huddy SM, Bethke S, Hendrikse M, Hitzeroth II, Rybicki EP. Production of human papillomavirus pseudovirions in plants and their use in pseudovirion-based neutralisation assays in mammalian cells. Sci Rep (2016) 6:20431. doi: 10.1038/srep20431

51. Leder C, Kleinschmidt JA, Wiethe C, Müller M. Enhancement of capsid gene expression: Preparing the human papillomavirus type 16 major structural gene L1 for DNA vaccination purposes. J Virol (2001) 75:9201–09. doi: 10.1128/JVI.75.19.9201-9209.2001

52. Lenzi P, Scotti N, Alagna F, Tornesello ML, Pompa A, Vitale A, De Stradis A, Monti L, Grillo S, Buonaguro FM, Maliga P, Cardi T. Translational fusion of chloroplast-expressed human papillomavirus type 16 L1 capsid protein enhances antigen accumulation in transplastomic tobacco. Transgen Res (2008) 17:1091– 1102. doi: 10.1007/s11248-008-9186-3

53. Lizotte PH, Wen AM, Sheen MR, Fields J, Rojanasopondist P, Steinmetz NF, Fiering S. *In situ* vaccination with Cowpea Mosaic Virus nanoparticles suppresses metastatic cancer. Nat Nanotech (2015) 11:295–303. doi: 10.1038/nnano.2015.292

54. Ljubojevic S, Skerlev M. HPV-associated diseases. Clin Derm (2014) 32:227-234. doi: 10.1016/j.clindermatol.2013.08.007

55. Lu B, Lim JM, Yu B, Song S, Neeli P, Sobhani N, K P, Bonam SR, Kurapati R, Zheng J, Chai D. The next-generation DNA vaccine platforms and delivery systems: advances, challenges and prospects. Front Immunol (2024) 15:1332939. doi: 10.3389/FIMMU.2024.1332939

56. Mach H, Volkin DB, Troutman RD, Wang B, Luo Z, Jansen KU, Shi L. Disassembly and reassembly of yeast-derived recombinant human papillomavirus virus-like particles (HPV VLPs). J Pharm Sci (2006) 95:2195– 2206. doi: org/10.1002/jps.20696

57. Maclean J, Koekemoer M, Olivier AJ, Stewart D, Hitzeroth II, Rademacher T, Fischer R, Williamson AL, Rybicki EP. Optimization of human papillomavirus type 16 (HPV-16) L1 expression in plants: comparison of the suitability of different HPV-16 L1 gene variants and different cell-compartment localization. J Gen Virol (2007) 88:1460–69. doi: org/10.1099/vir.0.82718-0

58. Marsian J, Lomonossoff GP. Molecular pharming – VLPs made in plants. Curr Op Biotech (2016) 37:201–206. doi: 10.1016/j.copbio.2015.12.007

59. Matic S, Rinaldi R, Masenga V, Noris E. Efficient production of chimeric Human papillomavirus 16 L1 protein bearing the M2e influenza epitope in *Nicotiana benthamiana* plants. BMC Biotech (2011) 11:106. doi: 10.1186/1472-6750-11-106

60. McCarthy MP, White WI, Palmer-Hill F, Koenig S, Suzich JA. Quantitative disassembly and reassembly of human papillomavirus type 11 virus like particles *in vitro*. J Virol (1998) 72:32–41. doi: 10.1128/JVI.72.1.32-41.1998

61. Michelfelder S, Parsons J, Bohlender LL, Hoernstein SNW, Niederkrüger H, Busch A, Krieghoff N, Koch J, Fode B, Schaaf A, Frischmuth T, Pohl M, Zipfel PF, Reski R, Decker EL, Häffner K. Moss-produced, glycosylation-optimized human factor H for therapeutic application in complement disorders. J Am Soc Nephrol (2017) 28:1462–74. doi: org/10.1681/ASN.2015070745

62. Michelfelder S, Fischer F, Wäldin A, Hörle KV, Pohl M, Parsons J, Reski R, Decker EL, Zipfel PF, Skerka C, Häffner K. The MFHR1 fusion protein is a novel synthetic multitarget complement inhibitor with therapeutic potential. J Am Soc Nephrol (2018) 29:1141–53. doi: 10.1681/ASN.2017070738

63. Milferstaedt SWL, Joest M, Bohlender LL, Hoernstein SNW, Özdemir B, Decker EL, van der Does C, Reski R. Differential GTP-dependent in-vitro polymerization of recombinant Physcomitrella FtsZ proteins. Sci Rep (2025) 15:3095. doi: 10.1038/s41598-024-85077-6

64. Mitsubishi Chemical Group Corporation. Overseas consolidated subsidiary, Medicago to cease operations (2023). https://www.mcgc.com/english/news_release/pdf/01468/01708.pdf [Accessed May 21, 2018].

65. Munoz C, Schröder K, Henes B, Hubert J, Leblond S, Poigny S, Reski R, Wandrey F. Phytochemical exploration of ceruchinol in moss: A multidisciplinary study in biotechnological cultivation of *Physcomitrium patens* (Hedw.) Mitt. Appl Sci (2024) 14:1274. doi: 10.3390/app14031274

66. Mori S, Ozaki S, Yasugi T, Yoshikawa H, Taketani Y, Kanda. Inhibitory cis-element-mediated decay of human papillomavirus type 16 L1-transcript in undifferentiated cells. Mol Cell Biochem (2006) 288:47–57. doi: 10.1007/s11010-006-9117-7

67. Mortola E, Roy P. Efficient assembly and release of SARS coronavirus-like particles by a heterologous expression system. FEBS Lett (2004) 576:174–178. doi: 10.1016/J.FEBSLET.2004.09.009

68. Murén E, Nilsson A, Ulfstedt M, Johansson M, Ronne H. Rescue and characterization of episomally replicating DNA from the moss *Physcomitrella*. Proc Nat Acad Sci USA (2009) 106:19444–49. doi: 10.1073/pnas.0908037106

69. Muthamilselvan T, Khan MRI, Hwang I. Assembly of human papillomavirus 16 L1 protein in *Nicotiana benthamiana* chloroplasts into highly immunogenic Virus-Like Particles. J Plant Biol (2023) 66:331–340. doi: 10.1007/s12374-023-09393-6

70. Naupu PN, van Zyl AR, Rybicki EP, Hitzeroth II. Immunogenicity of plant-produced human papillomavirus (HPV) virus-like particles (VLPs). Vacc (2020) 8:740. doi: 10.3390/VACCINES8040740

71. Niederkrüger H, Busch A, Dabrowska-Schlepp P, Krieghoff N, Schaaf A, Frischmuth T. „Single-use processing as a safe and convenient way to develop and manufacture moss-derived biopharmaceuticals”. In: Eibl R, Eibl D, editors. Single-use technology in biopharmaceutical manufacture, 2nd edition. Hoboken NJ: Wiley (2019). p.311–318. 10.1002/9781119477891.ch28

72. Nooraei S, Bahrulolum H, Hoseini ZS, Katalani C, Hajizade A, Easton AJ, Ahmadian G. Virus-like particles: preparation, immunogenicity and their roles as nanovaccines and drug nanocarriers. J Nanobiotech (2021) 19:59. doi: 10.1186/s12951-021-00806-7

73. Orellana-Escobedo L, Rosales-Mendoza S, Romero-Maldona A, Parsons J, Decker EL, Monreal-Escalante E, Moreno-Fierros L, Reski R. An Env-derived multi-epitope HIV chimeric protein produced in the moss *Physcomitrella patens* is immunogenic in mice. Plant Cell Rep (2015) 34:425–433. doi: 10.1007/s00299-014-1720-6

74. Park MA, Kim HJ, Kim HJ. Optimum conditions for production and purification of human papillomavirus type 16 L1 protein from *Saccharomyces cerevisiae*. Prot Expr Pur (2008) 59:175–181. doi: 10.1016/j.pep.2008.01.021

75. Parsons J, Altmann F, Arrenberg CK, Koprivova A, Beike AK, Stemmer C, Gorr G, Reski R, Decker EL. Moss- based production of asialo-erythropoietin devoid of Lewis A and other plant-typical carbohydrate determinants. Plant Biotech J (2012) 10:851–861. doi: 10.1111/J.1467-7652.2012.00704.X

76. Peng W, Rayaprolu V, Parvate AD, Proner MF, Hui S, Parekh D, Shaffer K, Yu X, Saphire EO, Snijder J. Glycan shield of the ebolavirus envelop glycoprotein GP. Comm Biol (2022) 5:785. doi: 10.1038/s42003-022-03767-1

77. Pineo CB, Hitzeroth II, Rybicki EP. Immunogenic assessment of plant-produced human papillomavirus type 16 L1/L2 chimaeras. Plant Biotech J (2013) 11:964–975. 10.1111/pbi.12089

78. Ramezaniaghdam M, Bohlender LL, Parsons J, Hoernstein SNW, Decker EL, Reski R. Recombinant production of spider silk protein in Physcomitrella photobioreactors. Plant Cell Rep (2025) 44:103. doi: 10.1007/s00299-025-03485-y

79. Rempfer C, Wiedemann G, Schween G, Kerres KL, Lucht JM, Horres R, Decker EL, Reski R. Autopolyploidization affects transcript patterns and gene targeting frequencies in Physcomitrella. Plant Cell Rep (2022) 41:153–173. doi: 10.1007/s00299-021-02794-2

80. Reski R. Development, genetics and molecular biology of mosses. Bot Acta (1998) 111:1–15. doi: 10.1111/j.1438-8677.1998.tb00670.x

81. Reski R. Quantitative moss cell biology. Curr Op Plant Biol (2018) 46:39–47. doi: 10.1016/J.PBI.2018.07.005

82. Reski R, Abel WO. Induction of budding on chloronemata and caulonemata of the moss, *Physcomitrella patens*, using isopentenyladenine. Planta (1985) 165:354–358. doi: 1007/BF00392232

83. Reski R, Parsons J, Decker EL. Moss-made pharmaceuticals: from bench to bedside. Plant Biotech J (2015) 13:1191–98. doi: 10.1111/pbi.12401

84. Reski R, Bae H, Simonsen HT. *Physcomitrella patens*, a versatile synthetic biology chassis. Plant Cell Rep (2018) 37:1409–17. doi: 10.1007/S00299-018-2293-6

85. Roden RBS, Stern PL. Opportunities and challenges for human papillomavirus vaccination in cancer. Nat Rev Cancer (2018) 18:240–254. doi: 10.1038/nrc.2018.13

86. Rommel O, Dillner J, Fligge C, Bergsdorf C, Wang X, Selinka HC, Sapp M. Heparan sulfate proteoglycans interact exclusively with conformationally intact HPV L1 assemblies: Basis for a virus-like particle ELISA. J Med Virol (2005) 75:114–121. doi: 10.1002/jmv.20245

87. Rosales-Mendoza S, Orellana-Escobedo L, Romero-Maldonado A, Decker EL, Reski R. The potential of *Physcomitrella patens* as a platform for the production of plant-based vaccines. Exp Rev Vacc (2014) 13:203–212. doi: 10.1586/14760584.2014.872987

88. Ross TM, Mahmood K, Crevar CJ, Schneider-Ohrum K, Heaton PM, Bright RA. A trivalent virus-like particle vaccine elicits protective immune responses against seasonal influenza strains in mice and ferrets. PLOS ONE (2009) 4:e6032. doi: 10.1371/JOURNAL.PONE.0006032

89. Ruiz-Molina N, Parsons J, Müller M, Hoernstein SNW, Bohlender LL, Pumple S, Zipfel PF, Häffner K, Reski R, Decker EL. A synthetic protein as efficient multitarget regulator against complement over-activation. Comm Biol (2022a) 5:152. doi: 10.1038/s42003-022-03094-5

90. Ruiz-Molina N, Parsons J, Schroeder S, Posten C, Reski R, Decker EL. Process engineering of biopharmaceutical production in moss bioreactors *via* model-based description and evaluation of phytohormone impact. Front Bioeng Biotech (2022b) 10:837965. doi: 10.3389/fbioe.2022.837965

91. Rybicki EP. Plant molecular farming of virus-like nanoparticles as vaccines and reagents. WIREs Nanomed Nanobiotech (2020) 12:e1587. doi: 10.1002/wnan.1587

92. Sapp M, Fligge C, Petzak I, Robin Harris J, Streeck RE. Papillomavirus assembly requires trimerization of the major capsid protein by disulfides between two highly conserved cysteines. J Virol (1998) 72:6186–6189. doi: 10.1128/jvi.72.7.6186-6189.1998

93. Schmidt CQ, Slingsby FC, Richards A, Barlow PN. Production of biologically active complement factor H in therapeutically useful quantities. Prot Expr Pur (2011) 76:254–263. doi: 10.1016/J.PEP.2010.12.002

94. Schween G, Hohe A, Koprivova A, Reski R. Effects of nutrients, cell density and culture techniques on protoplast regeneration and early protonema development in a moss, Physcomitrella patens. J Plant Phys (2003) 160:209–212. doi: 10.1078/0176-1617-00855

95. Scotti N, Rybicki EP. Virus-like particles produced in plants as potential vaccines. Exp Rev Vacc (2013) 12:211–224. doi: 10.1586/erv.12.147

96. Serradell MC, Rupil LL, Martino RA, Prucca CG, Carranza PG, Saura A, Fernández EA, Gargantini PR, Tenaglia AH, Petiti JP, Tonelli RR, Reinoso-Vizcaino N, Echenique J, Berod L, Piaggio E, Bellier B, Sparwasser T, Klatzmann D, Luján HD (2019) Efficient oral vaccination by bioengineering virus-like particles with protozoan surface proteins. Nat Comm (2019) 10:361. doi: 10.1038/s41467-018-08265-9

97. Shen JS, Busch A, Day TS, Meng XL, Yu CI, Dabrowska-Schlepp P, Fode B, Niederkrüger H, Forni S, Chen S, Schiffmann R, Frischmuth T, Schaaf A. Mannose receptor-mediated delivery of moss-made α- galactosidase A efficiently corrects enzyme deficiency in Fabry mice. J Inher Metab Dis (2016) 39:293–303. Doi: 10.1007/S10545-015-9886-9

98. Shi L, Sanyal G, Ni A, Luo Z, Doshna S, Wang B, Graham TL, Wang N, Volkin DB. Stabilization of human papillomavirus virus-like particles by non-ionic surfactants. J Pharm Sci (2005) 94:1538–51. doi: 10.1002/jps.20377

99. Shi L, Sings HL, Bryan JT, Wang B, Wang Y, Mach H, Kosinski M, Washabaugh MW, Sitrin R, Barr E. GARDASIL®: Prophylactic human papillomavirus vaccine development – From bench top to bed-side. Clin Pharm Therap (2007) 81:259–264. doi: 10.1038/SJ.CLPT.6100055

100. Šmídková M, Müller M, Thönes N, Piuko K, Angelisová P, Velemínský J, Angelis KJ. Transient expression of human papillomavirus type 16 virus-like particles in tobacco and tomato using a tobacco rattle virus expression vector. Biol Plant (2010) 54:451–460. doi: 10.1007/s10535-010-0081-4

101. Song JM, Wang BZ, Park KM, van Rooijen N, Quan FS, Kim MC, Jin HT, Pekosz A, Compans RW, Kang SM. Influenza virus-like particles containing M2 induce broadly cross protective immunity. PLOS ONE (2011) 6:e14538. doi: 10.1371/JOURNAL.PONE.0014538

102. Studentsov YY, Schiffman M, Strickler HD, Ho GYF, Pang YYs, Schiller J, Herrero R, Burk RD. Enhanced enzyme-linked immunosorbent assay for detection of antibodies to virus-like particles of human papillomavirus. J Clin Microbiol (2002) 40:1755–60. doi: 10.1128/JCM.40.5.1755-1760.2002

103. Strepp R, Scholz S, Kruse S, Speth V, Reski R. Plant nuclear gene knockout reveals a role in plastid division for the homolog of the bacterial cell division protein FtsZ, an ancestral tubulin. Proc Nat Acad Sci USA (1998) 95:4368–73. doi: 10.1073/PNAS.95.8.4368

104. Top O, Parsons J, Bohlender LL, Michelfelder S, Kopp P, Busch-Steenberg C, Hoernstein SNW, Zipfel PF, Häffner K, Reski R, Decker EL. Recombinant production of MFHR1, a novel synthetic multitarget complement inhibitor, in moss bioreactors. Front Plant Sci (2019) 10:260. doi: 10.3389/FPLS.2019.00260

105. Top O, Milferstaedt SWL, van Gessel N, Hoernstein SNW, Özdemir B, Decker EL, Reski R. Expression of a human cDNA in moss results in spliced mRNAs and fragmentary protein isoforms. Comm Biol (2021) 4:964. doi: 10.1038/s42003-021-02486-3

106. Torresi J. The rationale for a preventative HCV virus-like particle (VLP) vaccine. Front Microbiol (2017) 8:2163. doi: 10.3389/fmicb.2017.02163

107. van Wijk KJ. Intra-chloroplast proteases: A holistic network view of chloroplast proteolysis. Plant Cell (2024) 36:3116–30. 10.1093/plcell/koae178

108. Varsani A, Williamson AL, Rose RC, Jaffer M, Rybicki EP. Expression of Human papillomavirus type 16 major capsid protein in transgenic *Nicotiana tabacum* cv. Xanthi. Arch Virol (2003) 148:1771–86. doi: 10.1007/s00705-003-0119-4

109. Weise A, Rodriguez-Franco M, Timm B, Hermann M, Link S, Jost W, Gorr G. Use of *Physcomitrella patens* actin 5′ regions for high transgene expression: importance of 5′ introns. Appl Microbiol Biotech (2006) 70:337–345. doi: 10.1007/s00253-005-0087-6

110. Welsch S, Müller B, Kräusslich HG. More than one door - Budding of enveloped viruses through cellular membranes. FEBS Lett (2007) 581:2089–97. doi: 10.1016/J.FEBSLET.2007.03.060

111. Wiedemann G, van Gessel N, Köchl F, Hunn L, Schulze K, Maloukh L, Nogué F, Decker EL, Hartung F, Reski R. RecQ helicases function in development, DNA repair, and gene targeting in *Physcomitrella patens*. Plant Cell (2018) 30:717–736. doi: 10.1105/tpc.17.00632

112. World Health Organization. Considerations for human papillomavirus (HPV) vaccine product choice (2024). https://iris.who.int/bitstream/handle/10665/376546/9789240089167-eng.pdf [Accessed May 21, 2025].

113. Wu M, Huang H, Tang Y, Ren X, Jiang X, Tian M, Li W. Unveiling the multifaceted realm of human papillomavirus: a comprehensive exploration of biology, interactions, and advances in cancer management. Front Immunol (2024) 15:1430544. doi: 10.3389/fimmu.2024.1430544

114. Zahin M, Joh J, Khanal S, Husk A, Mason H, Warzecha H, Ghim SJ, Miller DM, Matoba N, Jenson AB. Scalable production of HPV16 L1 protein and VLPs from tobacco leaves. PLOS ONE (2016) 11:e0160995. doi: 10.1371/JOURNAL.PONE.0160995

115. Zhang S, Yong LK, Li D, Cubas R, Chen C, Yao Q. Mesothelin virus-like particle immunization controls pancreatic cancer growth through CD8+ T cell induction and reduction in the frequency of CD4+ foxp3+ ICOS- regulatory T cells. PLOS ONE (2013) 8:e68303 doi: 10.1371/JOURNAL.PONE.0068303

116. Zhao X, Schwartz S. Inhibition of HPV-16 L1 expression from L1 cDNAs correlates with the presence of hnRNP A1 binding sites in the L1 coding region. Virus Genes (2008) 36:45–53. doi: 10.1007/s11262-007-0174-0

117. Zlotnick A, Mukhopadhyay S. Virus assembly, allostery and antivirals. Trends Microbiol (2011) 19:14–23.doi: 10.1016/J.TIM.2010.11.003

118. zur Hausen H (2002) Papillomaviruses and cancer: From basic studies to clinical application. Nat Rev Cancer (2002) 2:342–350. doi: 10.1038/nrc798

119. zur Hausen H, de Villiers EM, Gissmann L. Papillomavirus infections and human genital cancer. Gyn Oncol (1081) 12:S124–S128. doi 10.1016/0090-8258(81)90067-6

